# Maternal experience unmasks temporal integration for vocalization processing

**DOI:** 10.1101/2023.09.13.557594

**Authors:** Amy M. LeMessurier, Gurket Kaur, Julia I. Paraiso, Ayat A. Agha, Janaye Stephens, Robert C. Froemke

## Abstract

New parents become highly responsive to infant cries in parallel with auditory cortical plasticity enhancing representations of these calls. To promote caretaking behavior, increased cortical activity must be appropriately routed to downstream areas in combination with suppression of circuits that could promote aversive responses to infant cries. We investigated responses to multi- syllable mouse pup calls in cortical output neurons projecting to either the auditory striatum or auditory midbrain (inferior colliculus) using longitudinal 2-photon imaging and in vivo optically- tagged whole-cell recordings. In mice with maternal experience, corticocollicular but not corticostriatal projections responded to pup calls over extended durations and were required for pup retrieval behavior, indicating that corticocollicular output neurons integrated multiple call syllables. The two populations diverged with parental experience, enhancing sustained responses in corticocollicular projections and reducing responses in corticostriatal projections. We performed simultaneous Neuropixels recordings from auditory cortex, thalamus, and midbrain to examine the temporal structure of call responses across areas, and to determine the influences of feedforward and feedback processing. Facilitation of later syllables increased in midbrain of maternally- experienced animals, together with reduced firing rates in a subset of units that were weakly coupled to cortical and midbrain activity. Overall, maternal experience leads to broad suppression of subcortical excitation in response to pup calls while selectively unmasking a subset of midbrain neurons that integrate over multiple syllables.

## Introduction

Perception of vocalizations is one of the most important functions of the auditory system in social animals. Vocalizations made by infants to indicate distress are common in altricial species, and survival depends on care-takers accurately identifying and responding to these calls^1–3^. Young mouse pups emit a specific category of ultrasonic vocalizations (USVs) when they are isolated outside of the warmth of the nest. Experienced moms (dams) will respond to these calls by searching for and retrieving the vocalizing pup^4–7^. Retrieval can also be learned by virgins that have previously never interacted with pups by co-housing with a dam and litter^5–8^, and plasticity in left auditory cortex responses to USVs has been identified as a key correlate of this learning^6,8–13^. The temporal structures of USVs are particularly important for eliciting behavior^10,14,15^, and variations in syllable durations and repetition rates may enable perceptual and neural discrimination of distinct categories^16^. In contrast, pup USVs are aversive to non-maternal (naïve) nulliparous females, driving avoidance^10,17^. Changes in cortical representations with the onset of maternal behavior refine selectivity for temporal features, which may reflect the semantic nature of the calls^8–10^.

In addition to segmentation and categorization of distinct vocal cues, accurately responding to vocalizations requires communication to downstream areas to guide behavior^1^. In mammals, the central auditory pathway consists of both feedforward connections that progress from the ears to higher cortical areas, as well as descending cortical projections that innervate much of the feed- forward pathway, including brainstem^18,19^, midbrain^20,21^, and thalamus^22,23^. Synapses from these top-down projections are numerous^24^, and are key for predictive coding computations^25–27^ that may support vocal attention and speech comprehension^28^. In humans, the role of top-down modulation in predictive coding has primarily been studied within the cortex^25^, but evidence from animal studies suggests that corticofugal projections to early areas in the auditory system can selectively process behaviorally-important sounds while suppressing noise, to drive representational plasticity enhancing discrimination/detection of the most important cues^29,30^. Neurons in layer 5 of auditory cortex send corticofugal projections to areas earlier in the auditory hierarchy, as well as feedforward projections to the striatum and amygdala^31^. Corticostriatal projections from auditory cortex primarily innervate the posterior striatum^32^, and facilitation of these synapses has been correlated with learning of trained acoustic cues in auditory perceptual learning tasks^33,34,35^. As the striatum is known to be involved in action selection and motor sequencing^36–40^, we initially hypothesized that this projection could serve as a direct link between auditory perception of pup vocalizations and motor planning and execution of retrieval.

Corticofugal projections provide a means for auditory cortex to modulate activity in earlier steps of the central auditory pathway. These projections are involved in both trained and innate auditory-guided behaviors, for example, re-learning to localize sounds following partial hearing loss^29^, appetitive perceptual training^41–43^, and flight responses to looming sounds^44^. We hypothesized that auditory cortex could facilitate pup retrieval by modulating USV representations in neurons that project more directly to motor execution circuits, or by suppressing activity in circuits that mediate pup avoidance. IC is the first area of the auditory pathway in which neurons extensively integrate acoustic input over time^45^, and which respond to USVs in multiple species^46–49^. Therefore, we also targeted projections from auditory cortex to IC. Many of these neurons may also send collateral axons to the thalamus, as well as posterior striatum and amygdala^18,31,50–53^, and could strongly modulate USV representations in multiple downstream areas simultaneously.

## Results

We first measured USV encoding in populations of neurons projecting to posterior striatum and IC using in vivo 2-photon calcium imaging, to ask how distinct subcortical circuits downstream of auditory cortex are specifically important for perception of USVs. We expressed jGCaMP7f or 8f in CC neurons (N=9 imaging fields from 8 female C57BL/7 mice) or CS neurons (N=10 fields from 6 females) by injecting rgAAV-hSyn-jGCaMP7/8f in either left IC or left posterior striatum (**Figure 1a**, left). During the same surgeries, chronic cranial windows were implanted over left auditory cortex and steel headbars were attached to the skull to enable head stabilization during imaging. Imaging experiments measuring USV and tone-evoked activity were performed 3-5 weeks following surgery. Following imaging experiments, brains were sectioned and imaged to visualize injection targeting and expression in auditory cortex. In all CC and CS mice, expression in cell bodies was most dense several hundred µm from the pial surface (**Figure 1a**, right). The bulk of labeled neurons had morphology matching that of L5 excitatory neurons, with large apical dendrites and high branching density in superficial layers of cortex.

**Figure 1.**
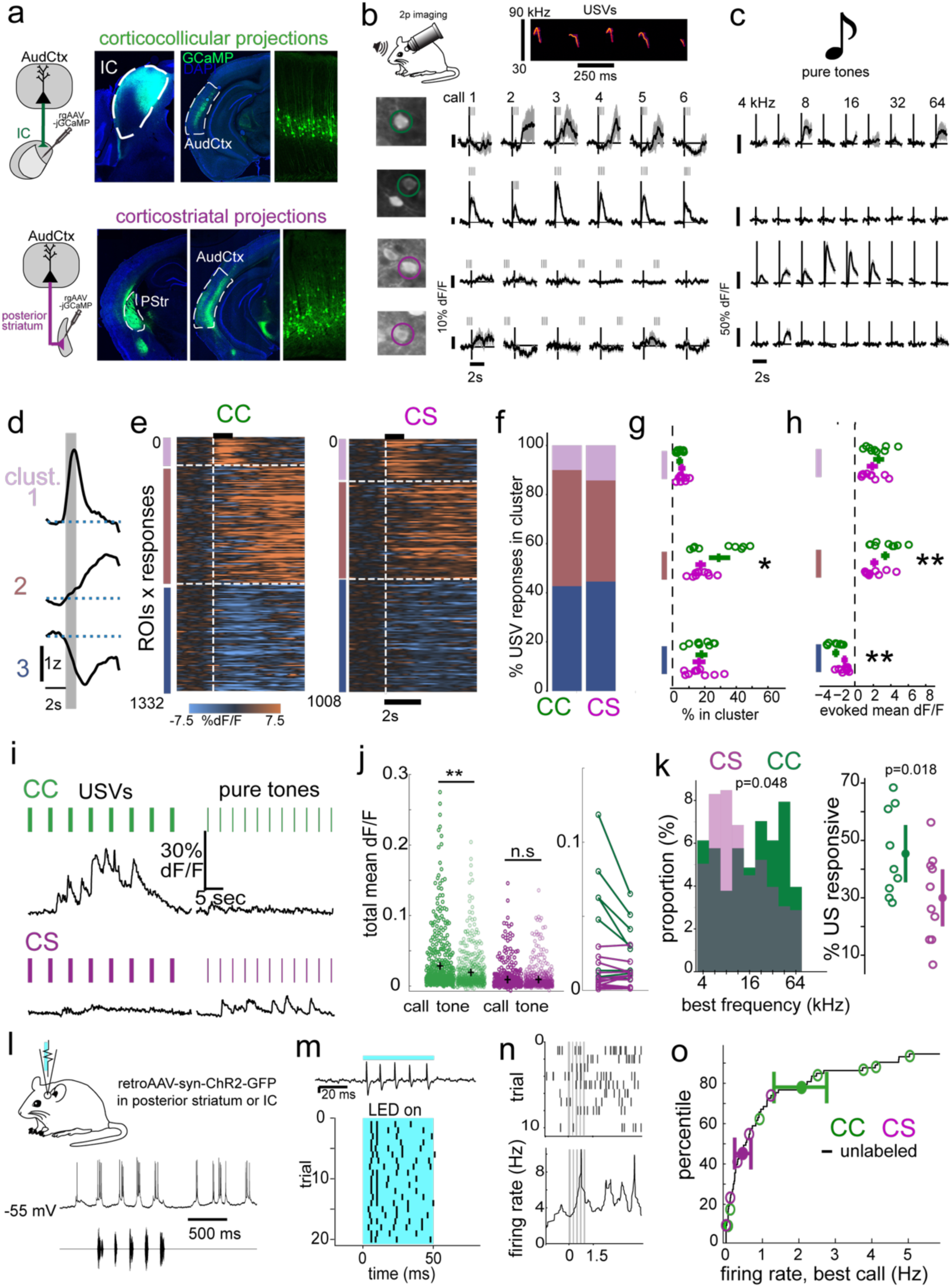
Corticocollicular projections respond to pup distress USVs over extended durations. **a**, Expression of jGCaMP7f or 8f in auditory corticocollicular (top, CC, green) or corticostriatal (bottom, CS, purple) projection neurons. Shown are histological images confirming injection targeting in fixed coronal sections from example brains following 2p imaging experiments. Fluorescence images of injection site, expression in auditory cortex, and high-magnification image of jGCaMP-expressing neurons in auditory cortex, respectively. IC: inferior colliculus, PStr: posterior striatum; AudCtx: auditory cortex. **b**, Responses to USVs during in vivo 2-photon imaging in awake head-fixed nulliparous females. Mean USV-evoked dF/F traces (average of 10-20 repeats/stimulus) shown relative to timing of USV playback for four example neurons/neurons depicted in left images. Top two rows: CC neurons; bottom rows: CS neurons. Black lines, means; gray shading, SEM; vertical black lines, stimulus onset; gray vertical hashmarks, individual USV syllables. **c**, Pure tone responses (half-octave spaced tones from 4-64 kHz). **d-h**, Clustered trial-averaged responses to USVs; there were three clusters representing onset responses (cluster 1), delayed responses (cluster 2), and suppression (cluster 3). **d**, Means of all significant evoked responses in each cluster. Grey bar, mean duration of all 6 USVs played. **e**, All trial-averaged evoked responses sorted by cluster; x-axis is time relative to USV onset, y-axis positions are individual USV responses from single neurons. Color, dF/F. **f**, Percent of significant evoked responses in each cluster from CC and CS neurons pooled across mice. Percentages do not include un-clustered or non-significant responses. **g**, Percentages of responses in each cluster for each individual animal (CC, N=9 imaging fields from 8 mice, green; CS, N=10 fields from 6 mice, purple; OLS regression, p=0.010 comparing cluster 2 CC vs CS). Horizontal lines, means and 95% confidence intervals (C.I.s). **h**, Mean call-evoked dF/F over 5 seconds following stimulus onset sorted by cluster (linear mixed-effects model with random intercepts for mouse identity; β: −0.55, p=0.054 comparing CS to CC; β: −0.69, p*<*0.001 comparing cluster 2 CC vs CS; β: 1.22, p*<*0.001 comparing cluster 3 CC vs CS). Circles, each mouse; lines, mean and 95% C.I. for responses across mice. All data collected on day 3 of cohousing or when animals displayed stable expert performance on pup retrieval assay. **i**, dF/F traces from example neurons imaged during epochs of call presentations (left) and tone presentations (right). Upper row: one corticocollicular neuron; lower row: one corticostriatal. Vertical bars denote timing of stimulus delivery, with inter-stimulus interval of 5 seconds for USVs and 3 seconds for tones. **j,** Total mean dF/F in all ROIs from N=9 corticocollicular labeled imaging fields from 8 mice (green, left) and N=10 corticostriatal fields from 6 mice averaged over entire blocks of USV or tone presentations (purple, right). Each circle is one cell’s mean; black crosses indicate population means. Right: total mean dF/F during tone and call blocks averaged within mouse. Each line depicts on mouse. Data were collected on day 3 of cohousing or when animals displayed stable expert performance. (Linear mixed-effects model with random intercepts per imaging field, β: 0.009, p=7.14*10^-10^ comparing calls vs tones in CC; β: 0.0006, p=0.68 comparing calls vs tones in CS; β: 0.031, p=0.002 effect of calls x projection). **k,** Left, distributions of best frequency for USV-responsive neurons, defined as the frequency evoking the largest trial-averaged dF/F in 2 seconds following stimulus onset for each neuron. Green is CC neurons, purple is CS. (Linear mixed-effects model, log-transformed best tone in USV-responsive neurons with random intercepts per imaging field; β: 13.9, p=0.048 comparing CC vs CS.) Right: percentages of neurons from each mouse that were significantly responsive to at least one tone in the ultrasonic range, including 32, 45, or 64 kHz. (Weighted Least Squares linear regression, proportion per imaging field of USV-responsive neurons that were responsive to tones in the US range; β: 0.17, p=0.018 comparing CC vs CS). **l-o**, Opto-tagged in vivo whole-cell and cell-attached recordings from CS and CC neurons. **l**, Example in vivo current-clamp recording from L5 auditory cortex neuron in awake mouse showing responses to USV syllables. **m**, Example opto-tagged CS neuron. **n**, Example CC raster and average spiking response evoked by best call (USV that evoked highest firing rate for that cell). **o**, Cumulative distribution of best call-evoked firing rate for putative L5 neurons recorded in current-clamp or cell-attached mode. Black line: all neurons including non-tagged (n=77 neurons from 21 retrieving females; green, n=8 CC neurons; purple, n=5 CS recordings).

We first examined representations of USVs and tones in expert retrieving nulliparous females (naïve virgins) cohoused with an experienced dam and her litter for several days. At least one day following habituation to head-fixation, awake mice were stabilized under a 2-photon microscope, and auditory stimuli were played through an ultrasonic speaker as we imaged left auditory cortex. We measured responses to a set of six pup isolation USVs (**Figures 1b, Extended Data Fig. 1a**), each comprised of 4-5 individual syllables and of durations ∼1 second, as well 500 ms pure tone pips at half-octave frequency intervals across the mouse hearing spectrum (4 to 64kHz, **Figure 1c**).

We examined USV and tone responses in CC vs CS neurons (**Figure 1b,c**). While there was a general diversity of response types across both populations (including neurons responsive to calls but not tones or vice versa), many neurons were reliably activated by USVs at long delays up to several seconds post stimulus onset. This suggested that neurons in both groups respond to specific features of calls, such as onsets and offsets, distinct motifs within calls, or integrated responses across multiple motifs. We sorted trial-averaged CC and CS responses into clusters following dimensionality reduction of all significant responses across mice (1444 USV responses from 488 CC neurons, 1119 USV responses from 428 CS neurons, **Extended Data Fig. S1a,b**). This yielded three groups of responses with distinct temporal profiles (**Figure 1d,e**). Both populations had a number of neurons with onset transients (cluster 1), but the predominant response had delayed positive transients that increased post USV-onset and remained elevated for at least 5 seconds post- USV onset (cluster 2). More CC than CS neurons were in this cluster (**Figure 1f,g**; p=0.02) and magnitudes were larger in CC than CS (**Figure 1h**; p=0.007). A large proportion of CC and CS neurons were also suppressed after USV onset (cluster 3).

Across mice, fractions of neurons in onset (cluster 1) or suppressed (cluster 3) groups were similar (**Figure 1g**), though suppression magnitude was greater in CC (**Figure 1h**, p=0.001). Overall, the strongest contrast between the two projections was the greater magnitudes and proportions of delayed responses in CC neurons, which could reflect increased temporal integration across multiple syllables. As temporal structures of USVs (e.g., durations and inter-syllable intervals) differ across categories, increased temporal integration could reflect a mechanism for categorical perception. In contrast to USV clusters, responses to pure tones were similar between CC and CS neurons, and broke out into either activated or suppressed (**Extended Data Fig. 1c-g**).

To confirm that our findings were not an artifact of clustering or oversampling from subsets of neurons responsive to multiple calls, we also compared three measures of call response magnitudes for each neuron that responded significantly to at least one call (**Extended Data Fig. 1h,i**): 1) early responses (0-2 sec post-stimulus onset) to the best USV and best frequency for each neuron; 2) later responses (2-5 sec post-onset) for calls and tones; and 3) most suppressed response-evoking calls and tones. For best late and most suppressed calls, CC neurons had larger responses to calls than CS neurons (late, β: 3.73, p=0.01; suppressed, β: -1.04, p=0.004). Late and suppressed tone responses were also larger in magnitude in CC neurons (late, β: 1.10, p=0.01; suppressed, β: -0.94, p<0.001); however, early responses were not significantly different for either calls or tones.

In addition to the differences we observed in USV-evoked and tone-evoked responses between CC and CS neurons, we observed a striking increase in activity in CC neurons during blocks of USV presentation beyond typical stimulus-evoked durations (**Figure 1 i-j**). We did not observe similar levels of increased long-duration firing rates in CC neurons during blocks of tones (CC mean dF/F over USV blocks vs. tone blocks: β=0.009, p<0.0001). In CS neurons, overall activity levels during USV blocks were lower than in CC (β=-0.03, p=0.002), and were equivalent during blocks of USVs and tones (β=-0.0007, p=0.68), suggesting that excitability in CC neurons selectively increases during barrages of pup USVs.

Previous studies examining the relationship between spectral tuning and USV-responsiveness in mature auditory cortex have found no correlation between USV-responsiveness and tuning for pure tones in the ultrasonic range^8,10,54,55^, though stronger correlations have been found early in development^56^ and in rats^12,57,58^. We quantified the best frequency out of 9 half-octave spaced tones for USV-responsive CC and CS neurons (**Figure 1k**). Neurons in both groups were tuned for tones across the mouse hearing range, but the distribution of CC neurons was shifted toward higher frequencies compared to CS (**Figure 1k** **left,** β = 3.40, p=0.048). Proportions of USV-responsive units with best frequencies in the ultrasonic range were also higher in CC-labeled mice compared to CS (**Figure 1k** **right**, β = 17.4%, p=0.018). Thus, USV-responsiveness may be driven by different synaptic inputs and circuit mechanisms across the two projections.

We next performed in vivo whole-cell and cell-attached recording from auditory cortical neurons including optogenetically-tagged CC and CS projection neurons. This was to ensure that differences in CC vs CS responses were not due to technical issues related to fluorescent imaging or Ca^2+^ indicator properties. We identified recorded neurons based on projection target by expressing channelrhodopsin in either projection and testing responsiveness to blue light^59^, and played calls (**Figure 1l-o**) and pure tones (**Extended Data Fig. 1j**). Current-clamp or cell-attached recordings were obtained from n=8 CC neurons from N=7 females (all virgins), and n=5 CS neurons from N=5 females (one dam and four virgins), as well as n=73 additional non-photo-tagged unidentified neurons. USV-evoked and tone-evoked firing rates varied greatly across these cells, ranging from the 90^th^ to 10^th^ percentiles of USVs responses and 55^th^ to 5^th^ percentiles of tone responses. However, mean firing rates for USVs were higher in CC vs CS neurons (**Figure 1o**, p=0.03, Student’s t-test) but not for tones (**Extended Data Fig. 1j,** p=0.09), consistent with our observations from imaging.

## Suppressing CC but not CS projections impaired pup retrieval

To determine whether subcortically-projecting neurons in auditory cortex were required for pup retrieval, we assayed retrieval performance during chemogenetic suppression of layer 5 neurons (**Figure 2**). We virally expressed inhibitory DREADD receptors (AAV1-DIO-hSyn-HM4D(gi)) in left auditory cortex of N=6 virgin female RBP4-Cre mice (GENSAT, UC-Davis) to selectively target layer 5 excitatory neurons. Retrieval performance was tested at least 2 weeks following surgery to allow for sufficient expression of the DREADDs construct. Mice were co-housed with wildtype (C57BL6j) dams with new litters beginning at postnatal day 0-1, and were cohoused for at least 3 days prior to pup retrieval testing to become alloparents, i.e., experienced retrievers. After confirming that co-housed virgins became expert retrievers (retrieving pups on ≥80% of trials), we tested pup retrieval performance after i.p. administration of CNO or saline.

**Figure 2.**
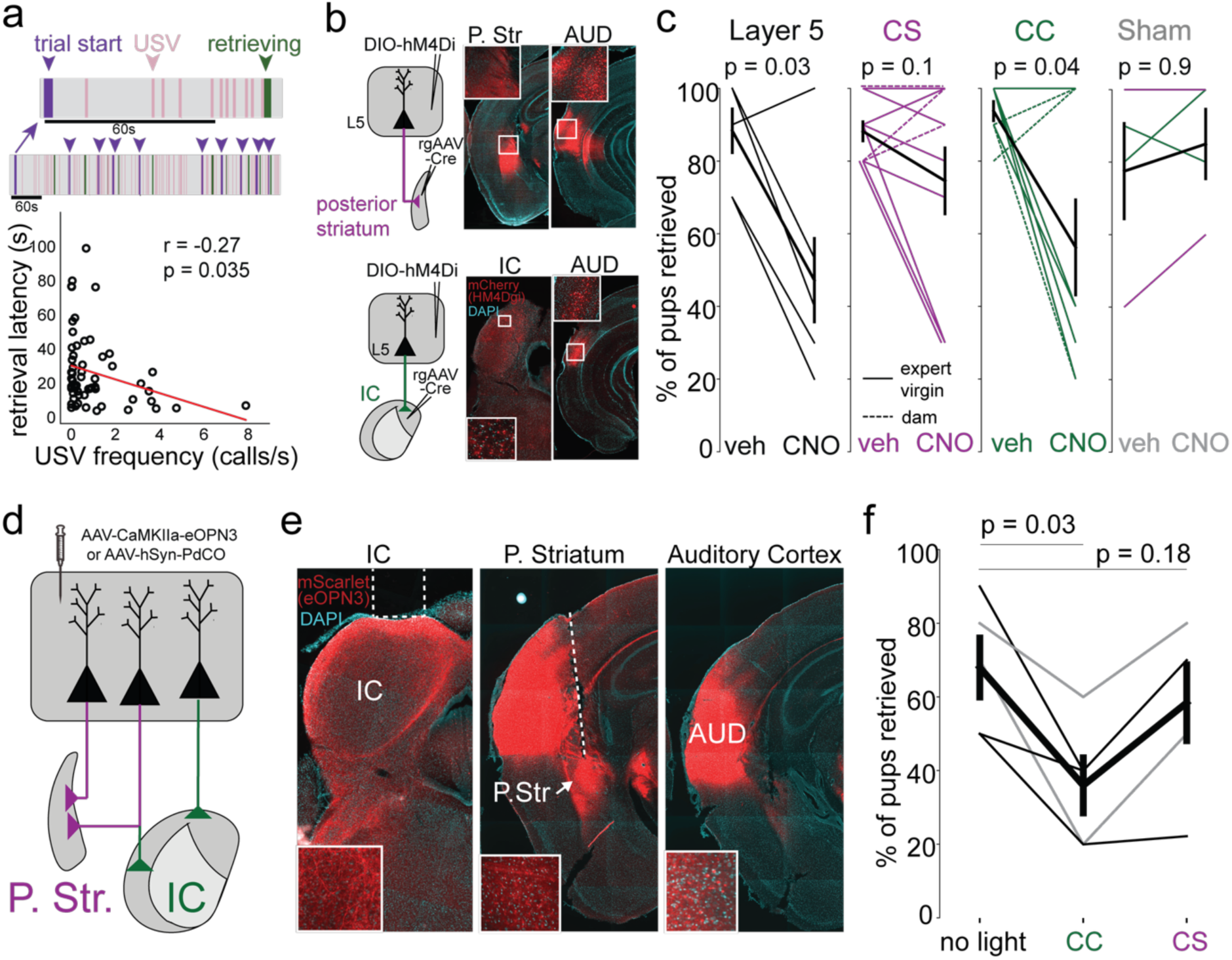
Chemo- and optogenetic suppression of left auditory cortex projections to IC impair pup retrieval. a, Top, example ethogram of 10 pup retrieval trials from a single mouse. Lower panel, entire duration; upper panel, zoom in of the first trial. Purple denotes trial start, defined as the time of experimenter placing pup in the arena. Pink lines indicate a bout of calls isolated by at least 1 second of silence. Green indicates time of retrieval, from the time the pup is picked up until dropped in or near the nest. Purple arrows in lower panel indicate trial starts. Bottom, negative correlation of frequency of USV emission (individual USVs per second) during a trial with the latency to retrieve on that trial. N=2 CS virgins and 4 CC virgins, all successful trials from saline block included in calculation of correlation. R=-0.27, p=0.035, Pearson’s R. **b**, Viral injection strategy and histological confirmation of targeting of CS (top) and CC (bottom) auditory cortex neurons. AAVrg-EF1a-Cre was injected into posterior striatum (P. Str.) or inferior colliculus (IC) using stereotactic coordinates for targeting. AAV1-hSyn-DIO-hM4D(Gi)-mCherry was injected into auditory cortex with targeting relative to anatomical landmarks (cranial sutures and blood vessels). Top left, example histological confirmation of injection targeting to posterior striatum (P. Striatum) and expression in auditory cortex. Bottom left, example confirmation of IC targeting. Middle, epifluorescence images of coronal sections containing posterior striatum (top) or IC (bottom). Right, auditory cortex. DAPI staining depicted in cyan, mCherry in red. Insets: higher magnification images of areas within white squares showing mCherry expression in fibers and cell bodies. **c**, Fraction of pups retrieved under each condition. Thin lines, individual animals (dashed line, dams; continuous line, virgins). Thick lines, means±SEM. Layer 5 suppression in RBP4-Cre mice with AAV1-hSyn-DIO-hM4D(gi)-mCherry in left auditory cortex impaired retrieval (N=6 experienced virgins; p=0.03, Wilcoxon rank sum test). Suppression of CS neurons in left auditory cortex did not impair retrieval (N=9, 3 dams, 6 experienced virgins; p=0.08). Suppression of CC neurons in left auditory cortex impaired retrieval (N=9, 3 dams, 6 experienced virgins; p=0.03). Sham mCherry-expressing controls were unimpaired after CNO (N=2 experienced virgins targeting CC, N=2 experienced virgins targeting CS, p=0.9). **d-e**, viral injection strategy and histological confirmation of expression and optic fiber targeting for optogenetic suppression of CC and CS terminals independently during pup retrieval. AAV-CaMKIIa-eOPN3-mScarlet-WPRE or AAV-hSyn-PdCO-eGFP-WPRE was injected into left auditory cortex as described for **b**. Dashed white lines in **e** indicate placement of optic fibers. Insets: higher magnification details of fibers and cell bodies in each labeled area. **f**, Fraction of pups retrieved under each condition. Thin lines, individual animals (gray, mice expressing PdCO; black, eOPN3). Thick line, means±SEM. Suppression of CC terminals in left IC impaired retrieval compared to no-light condition (N=5 experienced virgins; p=0.03, Wilcoxon rank sum test) but suppression of CS terminals in P. Striatum did not (p=0.18, Wilcoxon rank sum test).

To test retrieval, mice were habituated in the testing arena with 2-3 pups situated in a nest in the corner of the arena for 30+ minutes, following which single pups were introduced into a corner of the cage one at a time. On each of 10 trials mice were given 2 minutes to retrieve a pup to the nest. Retrieval took place under dim red light to reduce reliance on visual cues. Successful retrieval trials often occurred after the onset of USV emission by the displaced pup, and we found that the frequency of USVs emitted by the pup predicted faster retrievals by the dam, i.e., latency to retrieve pups was negatively correlated with frequency of USV emission during the trial (**Figure 2a**; N=5 expert retrieving virgins, r: -0.27, p=0.035). After saline injections, RBP4-Cre experienced virgins retrieved pups on 88±6% of trials, whereas on CNO trials, percent of pups retrieved decreased by roughly half to 47±12% (N=6, p=0.03, Wilcoxon rank sum test), indicating that pup retrieval depends on activity in layer 5 neurons from left auditory cortex.

Suppressing activity in CS neurons alone did not affect retrieval performance (**Figure 2b,c**). To selectively target CS neurons, Cre was retrogradely expressed by injecting rgAAV-Cre into the left posterior striatum, and Cre-dependent HM4D(gi)-mCherry virus or just mCherry for sham controls was injected into left auditory cortex. Correct targeting of injections and spread of expression were determined by visualizing mCherry in post-hoc histology for each mouse (**Figure 2b**). Retrieval performance after CNO administration was unchanged relative to saline on average in 8 experienced virgins and 3 dams (retrieval probability: saline, 88±3%; CNO, 75±1%, N=11, p=0.1). Similarly, no effect was seen in sham-injected controls (**Figure 2b,c**; retrieval probability, all shams: saline, 76±13%; CNO, 85±10%, p=0.9, paired t-test; CS sham only: saline, 70±3%; CNO, 80±2%).

The lack of effect of silencing CS neurons alone suggested that a different subset of layer 5 neurons was responsible for the diminished performance we observed when silencing left AC layer 5 more broadly. Thus, we assayed pup retrieval performance while suppressing activity in CC neurons using the same approach as described for CS neurons, with rgAAV-Cre injected into left auditory midbrain (**Figure 2b**). CNO injection to suppress CC neuron activity decreased pup retrieval (**Figure 2c**; retrieval probability: N=5 experienced virgins, 3 dams; saline, 94±3%; CNO, 56±13%, p=0.04). No effect was seen in sham mCherry-injected controls (**Figure 2c**, CC sham only: saline, 85±5%; CNO, 90±10%). Thus, corticofugal projections to IC are critical for pup retrieval in experienced alloparents and dams.

A subset of layer 5 excitatory neurons projecting to IC also send collateral projections to auditory thalamus, striatum, and lateral amygdala^18,31,50–53^; thus the reduction in pup retrieval we observed by retrograde viral expression of DREADDs may not have been specific to suppression of synaptic transmission at corticocollicular synapses. To independently suppress transmission at corticocollicular and corticostriatal synapses, we virally targeted expression of inhibitory optogenetic constructs that act at presynaptic terminals (eOPN3^60^ or PdCO^61^) to left auditory cortex (**Figure 2d-f**). We chronically implanted optic fibers in left IC and posterior striatum (N=3 nulliparous females injected with eOPN3 and N=2 injected with PdCO) and tested pup retrieval after allowing 4 weeks for viral expression and at least 3 days of cohousing with a dam and young pups. For each mouse we tested pup retrieval during delivery of 525 nm light (eOPN3) or 405 nm (PdCO) to IC and posterior striatum in separate blocks of 10 trials, as well as a no-light condition in which a fiber optic patch cable was attached but no light was delivered. The percent of pups retrieved was significantly lower with CC synapses suppressed compared to the no light condition (**Figure 2f**, no light: 68±8%; CC: 36±7%; N=5, p=0.03) but equivalent to the no-light condition with CS synapses suppressed (58±10%, p=0.18), further supporting the importance of CC projections over CS projections for pup retrieval.

## Plasticity of CC and CS projections with maternal experience

The necessity of CC but not CS projections for pup retrieval, and the enhanced USV responses in CC vs CS neurons together demonstrate that specializations in the auditory CC pathway are critical for maternal responses to pup calls. We wondered if USV representations in each pathway reflected innate sensitivity or were refined with parental experience. To test this, we performed 2- photon imaging of CC and CS neurons in virgins before and during co-housing with an experienced dam and litter. We measured responses to USVs and pure tones in virgins daily, starting before cohousing and for 2-4 days following (**Figure 3a,b**; N=4 virgins targeting CC projection neurons, N=3 virgins targeting CS neurons). Retrieval performance was tested daily after imaging. Pup retrieval generally improved over days of co-housing relative to day 0 (pre co- housing). In most mice, performance reached ≥80% after the third day of co-housing (**Figure 3c,e**).

**Figure 3.**
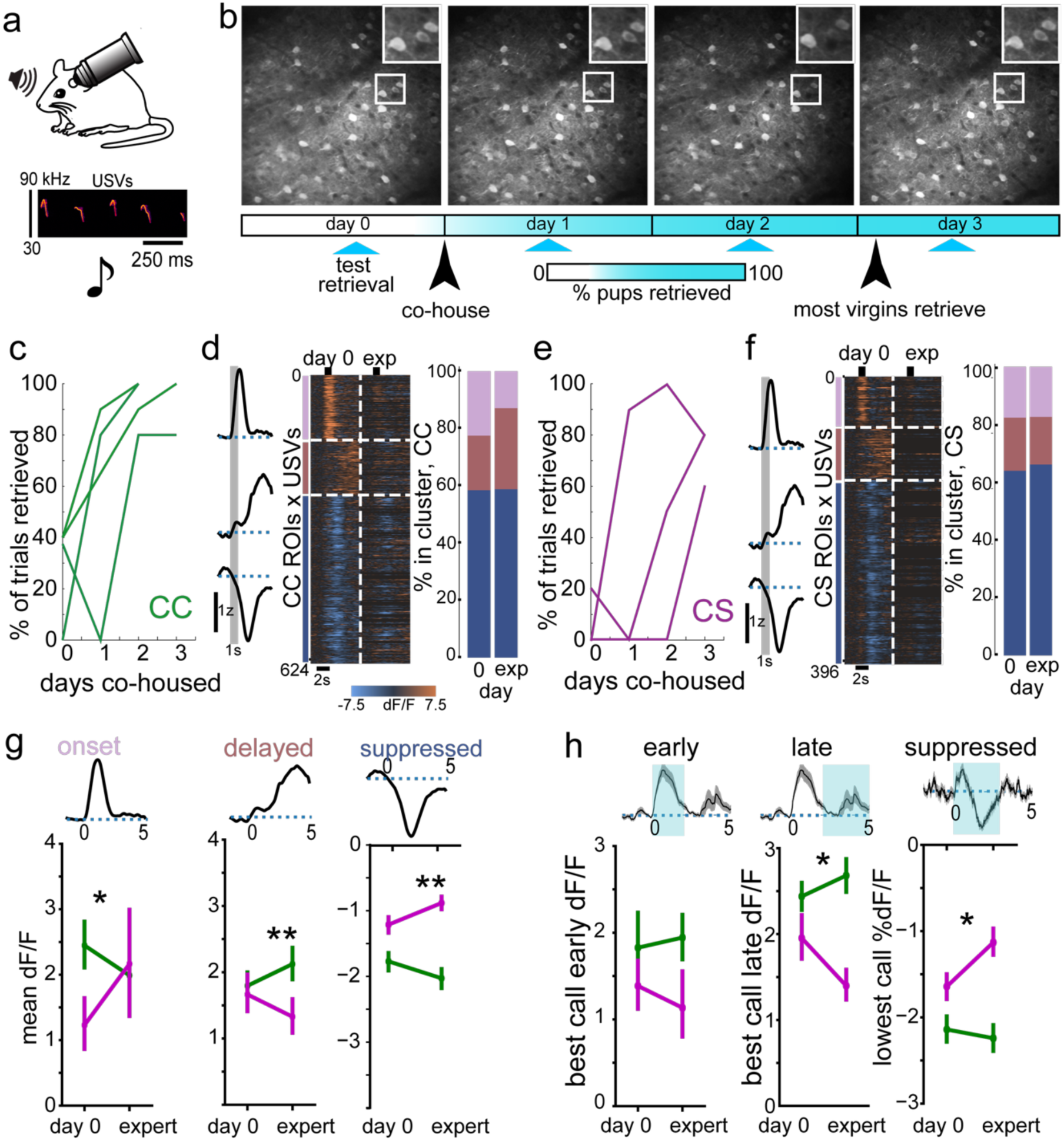
Longitudinal imaging during parental experience revealed divergent plasticity of CS vs CC connections. **a**, Experiment schematic. **b**, Example imaging field expressing jGCaMP8f on four consecutive imaging days. Images averaged over entire sessions (∼10^5^ frames). White squares, inset zoom-ins. Timeline shows image acquisition relative to co-housing. **c**, Pup retrieval in individual CC virgins over days co-housed with a dam and litter. Retrieval quantified on each day as percent of pups retrieved on all trials. **d-g**, Clustered trial-averaged responses to USVs. **d**, Left, means of all significant evoked responses in each cluster. Middle, trial-averaged evoked responses of CC neurons sorted by cluster on day 0, for all responses on day 0 and the first day of expert behavior. Only USV responses from neurons that were significant and clustered on day 0 are depicted. Color, dF/F. Black ticks, duration of USV presentation. Right, percent of significant evoked responses in each CC cluster across mice on day 0 and the first day of expert behavior. **f**, As **e** but for CS neurons. **g**, USV-evoked mean dF/F of CC (green) and CS (purple) neurons on day 0 and first day of expert retrieval for onset (day 0 projection effect, Z: -2.11, p=0.035, from OLS linear regression with SE clustered by mouse; projection by transition effect, Z: 1.52, p=0.129), delayed (CC day 0 vs expert effect, Z: 5.42, p<0.0001; CS day 0 vs expert effect, Z: - 2.65, p=0.008; projection by transition, Z: -4.11, p<0.0001), suppressed (CC day 0 vs expert effect, Z: -3.66, p<0.0001; CS day 0 vs expert effect, Z: 3.06, p=0.0022; projection by transition, Z: 4.35, p<0.0001). **h**, Left, early USV-evoked mean dF/F for best USV of each responsive neuron across mice within CC and CS groups. Early response is 0-2 seconds following call onset. Middle, best late USV-evoked mean dF/F; late response is 2-5 seconds post-call USV onset (best late response, Z: -2.63, p=0.0085, effect of transition day by projection; CS day 0 vs expert, Z: -2.29, p=0.022). Right, most-suppressed USV-evoked mean dF/F (most suppressed response, Z: 2.42, p=0.016, transition day by projection; CS day 0 vs expert, Z: 2.24, p=0.025).

USV-evoked dF/F magnitudes and tuning for individual calls were variable in tracked neurons, with both facilitated and suppressed responses observed each day for all stimuli. Despite individual neuron daily variability, cortical responsiveness to USVs was maintained at the population level, similar to previous findings of day-to-day variability in encoding of acoustic stimuli^62,63^. Most neurons that were detected on multiple days were not present on every imaging day, and many neurons with time-locked responses to specific calls on one day did not have the same responses or USV preference across all days. We clustered significant USV-evoked responses pooled across all days and mice. This included 1845 responses from 342 CC neurons (37.5% of CC responses) and 1424 responses from 256 CS neurons (38.4% of CS responses). Responses clustered into the same three groups based on temporal structures as in **Figure 1**, with all three clusters (early-activated, late-activated, suppressed) present in both CC and CS populations on all days of imaging (**Figure 3d,f**, **Extended Data Fig. 2**).

Since individual animal retrieval performance varied in terms of number of days of cohousing preceding expert retrieval, we compared responses to USVs and pure tones for each mouse between day 0 (before cohousing) and on the first day of expert retrieval (**Figure 3d,f,g**). The magnitudes of delayed and suppressed responses evolved in opposite directions for CC vs CS responses from day 0 to the first expert day (**Figure 3g**). In the delayed-response cluster, responses were initially similar between CC and CS on day 0 (β: -0.20, z: -0.89, p=0.374). However, CC- projecting neurons exhibited a significant increase in their delayed response on the first day of expert behavior compared to day 0 (β: 0.27, z: 5.42, p<0.001), while CS-projecting neurons exhibited a significant decrease in delayed response magnitude. Suppressed responses in CS neurons were initially smaller in magnitude than in CC neurons on day 0 (β: 0.52, z: 3.08, p=0.002), and magnitudes of suppression diverged further over the transition period with CC neurons becoming even more suppressed in experts (β: -0.21, z: -3.66, p<0.001), while CS neurons became less suppressed (post-hoc linear contrast: β: 0.36, z: 3.06, p=0.002). Fast onset responses (cluster 1) did not change in magnitude over the behavioral transition, although we confirmed that CC onset responses were somewhat stronger than for CS projections on day 0 (β: -1.06, z: -2.11, p=0.034). We observed similar trends when mean responses per cluster were quantified across chronological day (**Extended Data Figure 2d**). Together, these changes demonstrate the emergence of opposite population-level shifts in USV representations in CC vs CS projections in parallel with the transition to the alloparental state, likely resulting in opposing changes in call-evoked activity in downstream targets.

We also quantified best early, late, and suppressed responses to USVs on each day for every neuron that was imaged across at least two days (**Figure 3h**). We observed no projection-specific changes in early best call responses over the transition period (β: -0.33, z: -1.33, p=0.185), nor did we observe differences in either projection by chronological day (CC projections, β: 0.09, z: 0.50, p=0.619; CS projections, β: -0.24, z: -1.44, p=0.151). Best late response magnitudes were initially similar between CS and CC projections, but diverged significantly over the behavioral transition (β: -0.79, z: -2.73, p=0.006), primarily driven by a decrease in CS response magnitude (β: -0.55, z: -2.31, p=0.021). Similarly, suppressed call response magnitudes were not different on day 0 (β: 0.49, z: 1.53, p=0.127), but diverged significantly over the transition (β: 0.60, z: 2.43, p=0.015), due to decreased magnitude on the expert day in CS neurons (β: 0.50, z: 2.24, p=0.025). We observed similar effects when responses were quantified by day (**Extended Data Fig. 2e**). In agreement with the cluster-based quantifications (**Figure 3g, Extended Data Fig. 2d**), these findings support a model in which USV representations diverge in CC and CS populations over parental experience. Furthermore, long-term suppression of CS projections may be an important mechanism for the emergence of parental behavior in response to pup calls.

We also examined tone responses over the transition to alloparenting to ask whether plasticity in CC and CS neurons was specific to pup calls. Mean responses to tones clustered into a tone-excited cluster and a tone-suppressed cluster (**Extended Data Figure 2f-i**). On day 0, 64.3% of detectable neuron responses were excited and 35.7% were suppressed. In the activated cluster, contrary to what we observed for USV responses, neither projection changed in magnitude over experience (β: -1.23, z: -1.16, p=0.25), nor was there a difference between projections in excited response magnitude (β: -1.73, z: -1.46, p=0.14). In the suppressed cluster, CS responses decreased in magnitude over the transition (β: 2.28, z: 3.71, p=0.0002), in parallel with a decrease in the proportion of these responses. Consistent with these results, we observed no differences between projections or effects of transition day for early or late positive responses to each neuron’s best tone (p>0.05, all comparisons), but in the CS population, there was a reduction in magnitude in responses to the most activity-suppressing tone (β: 2.408, z: 2.13, p=0.033, **Extended Data Fig. 2j**). These findings were corroborated by mean tone responses per cluster averaged by chronological day (**Extended Data Fig. 2k-n**), though we did not observe any differences in most-suppressed tone responses averaged by chronological day (**Extended Data Fig. 2o**).

## Multisyllabic responses of single units to pup calls across the central auditory system

The preferential enhancement of delayed, activated responses in CC projection neurons with alloparenting experience indicates that the temporal structure of USV representations is plastic. Since the CC population included a higher proportion of these responses than CS, we reasoned that CC neurons may integrate acoustic input over longer durations, enabling recognition of timing features in vocalizations with repetitive structure. Therefore the transition to maternal behavior may require a shift toward encoding of multi-syllabic structure USVs in circuits driving pup retrieval. To investigate this hypothesis with higher resolution for timing features of USV responses, we conducted multi-area high-density Neuropixels recordings. We recorded simultaneously from auditory cortex, inferior colliculus (IC), and the medial geniculate body (MGB, the auditory thalamus) with Neuropixels probes in awake, head-fixed nulliparous females (N=11 naïve female virgins, 7 cohoused alloparents) during presentation of USVs and pure tones (**Figure 4a,b**). To test whether USV responses depended on temporal structure over multiple syllables rather than spectral cues present in individual syllables, we also played isolated individual syllables from one USV (call 3) spaced 2 seconds apart, in pseudorandom order.

**Figure 4.**
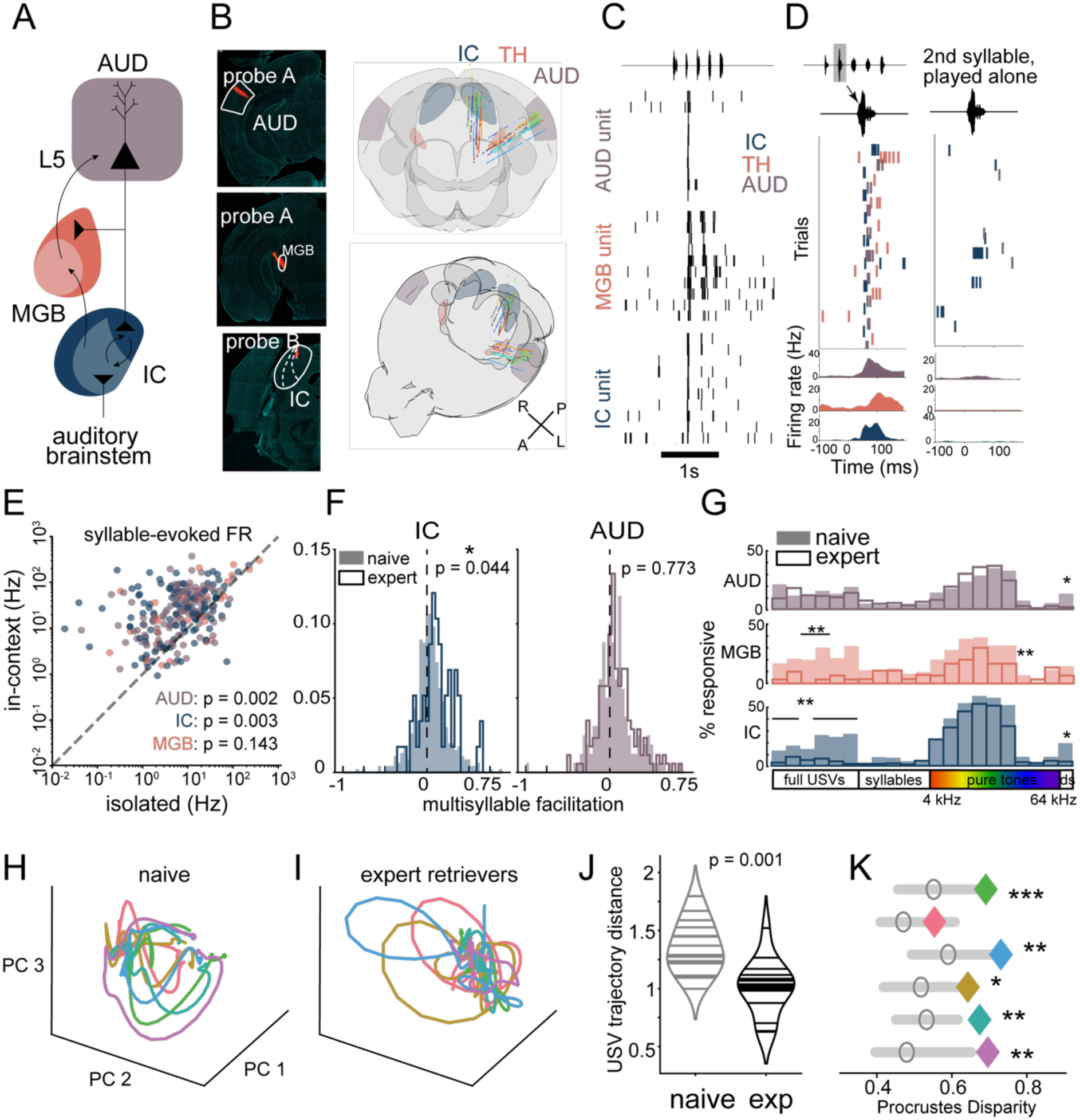
**Multisyllable infant USVs drive supralinear integration over sequential syllables in AUD, IC and MGB**. **a,** Schematic of inter-area connectivity within the central auditory system. **b,** Left, histological confirmation of silicon probe targeting in a primary auditory cortex (AUD), auditory thalamus (MGB) and inferior colliculus (IC) from a representative recording in an awake, head-fixed nulliparous female. Right, single unit locations in frontal (top) and rotated (bottom) view of a standard mouse brain, color-coded by mouse. Probe locations were reconstructed from diI tracks in histological sections. Shading denotes locations of auditory cortex (AUD), medial geniculate body (MGB), and inferior colliculus (IC). **c,** Spike rasters from three simultaneously recorded units in AUD, MGB, and IC that preferentially respond to the second syllable of an example USV. Top, waveform of multi-syllabic USV. **d,** Left, spike-timing and PSTHs relative to the second call syllable for the units depicted in **b**; right, responses to the second call syllable played in isolation. **e,** Scatter plot of isolated single USV syllable-evoked firing rates vs. firing rates to the identical syllables within a multi-syllable USV context. Each point represents the syllable evoking the largest-magnitude response for one USV-responsive single unit, color-coded by area. All values are baseline-subtracted relative to pre-stimulus period; negative values not shown but included in quantifications/statistics. n=65 AUD, 84 IC, 24 MGB units responsive to call 3. P-values as indicated obtained using Only Least Squares (OLS) linear models with cluster-robust standard error accounting for within-mouse variance**. f,** Distributions of multisyllable facilitation for each USV-responsive unit’s evoked firing rate to its preferred syllable relative to the first syllable in a multi-syllable USV. Data was pooled across mice within naïve (shaded) and expert retrieving (outlined) groups. P-vales denote comparisons between naïve and expert distributions within IC (left) and AUD (right) obtained from OLS linear regression with clustering by mouse. **g,** Percent of units pooled across recordings significantly driven by each acoustic stimulus, including 6 multisyllable USVs, 5 individual syllables from one USV, pure-tone pips, and trains of frequency-modulated sweeps. Top, AUD units; middle, MGB units, bottom IC units. For each bar plot, shaded bars indicate naïve and outlined indicate expert. Asterisks denote p<0.05 (*) and p<0.001 (**) comparing individual stimulus proportions. P-values were obtained using Generalized Estimating Equations accounting for intra-mouse correlation and corrected for multiple comparisons by False Discovery Rate method. **h,** Trial-averaged population trajectories for each exemplar USV projected into the first 3 principal components for visualization for all USV-responsive units pooled across naïve mice. Filled circles depict time of trajectory onset. **i,** As described in **h**, for expert-retrieving mice. **j,** Distributions of trajectory distances between each pair of USVs in naïve (gray) and expert (black) populations. P=0.0005, U=28, Mann-Whitney U test for distance by experience. **k**, Procrustes disparity assessing geometric divergence in over time for each exemplar USV between naïve and expert groups. Disparity was computed from trajectories in state-space defined by the number of principal components describing 80% of variance across units. Each colored diamond represents true disparity for one USV; gray shading indicates 0.05 to 0.95 confidence intervals for null distributions generated by permuting naïve and cohoused labels at mouse level and computing Procrustes Disparity for all possible permutations. Circles denote null mean. Asterisks: *, >95%; **>99%; ***, >99.9% of null distribution value. P-values per call: call 1: p=0.008, call 2: p=0.008, call 3: p=0.016, call 4: p=0.008, call 6: p<0.001.

We observed strongly USV-responsive units with evidence of temporal integration across syllables in all three regions (**Figure 4c-f**). Classic models of hierarchical feature encoding in sensory systems would predict that the complexity of features encoded in cortex would be greater than MGB and IC, and that the spatiotemporal structure of USV encoding across tonotopic maps would be correlated with spectral temporal tuning in individual neurons^64^. In contrast, we observed neurons selective for syllable position in all three regions. Furthermore, these cells were not responsive to the identical syllables played outside of the call context (**Figure 4c-e**). We compared single-syllable-evoked firing rates between individual syllables from one USV and its individual syllables played in isolation, using the peak response to preferred syllables for all units that responded significantly to the full USV (**Figure 4e**). In both cortex and IC, firing rates evoked by the best syllable for each unit were higher within the USV context then when played in isolation (cortex: median 30.7 vs. 2.1 Hz, Wilcoxon signed-rank test W: 332.0, p<0.0001, n=65 units; IC: median 26.5 vs. 0.5 Hz, W: 1016.0, p=0.0006, n=84 units). In contrast, peak syllable responses in MGB were not significantly modulated by acoustic context (median 12.4 Hz in-context vs. 5.6 Hz in isolation; Wilcoxon signed-rank W: 137.0, p=0.726, n=24 units). This indicates that individual syllable responses in both cortex and IC were strongly modulated by the surrounding syllables, which could be due to adaptation or integration of responses to preceding syllables.

We quantified facilitation of each USV-responsive unit’s maximum rate relative to the first syllable for the preferred USV (**Figure 4f**). In each area, later-syllable responses had variable facilitation ranging from -1.0 (i.e., responding only to first syllable) to 0.75 (much stronger responses to later syllable); distributions were similar across areas (auditory cortex mean multisyllable facilitation: 0.08±0.01, n=332 units, N=14 mice; IC: 0.05±0.01, n=517 units, N=15 mice; MGB: 0.04±0.02, n=203 units, N=9 mice). Units from experienced retrievers were more facilitated than in naïve mice in IC (naive: mean 0.06±0.02, n=459 units, N=9 virgins; experienced: 0.12±0.03, n=58 units, N=6 alloparents) but not auditory cortex (naïve: 0.06±0.02, cohoused: 0.09±0.02, β: 0.016, 95% CI=[-0.093, 0.125], p=0.773). Very few MGB units recorded in cohoused mice were USV-responsive (n=8 units from N=2 mice) so we were unable to accurately assess the effect of cohousing on adaptation in MGB USV responses. Overall, units in cortex and IC, but not MGB, were strongly modulated by multisyllabic context, with an increase in facilitation over multiple syllables in IC with alloparenting experience.

In addition to changing temporal integration of multiple USV syllables, maternal experience could increase sensitivity to pup calls by altering the overall proportions of USV-responsive neurons. Therefore we compared the fractions of significantly responding units in each area across stimuli between naïve and cohoused mice (**Figure 4g**). In auditory cortex, the overall proportions of responsive neurons were equivalent for all but one stimulus (n=243±32 units from N=4-7 expert mice; n=224±34 units from N=7-10 naïve mice). Out of the 21 acoustic stimuli tested, only 80 to 40 kHz sweeps repeated at typical USV inter-syllable interval were different, dropping from 13.0% in naive mice to 1.7% in alloparents (FDR-adjusted p=0.008). No pure tones or vocalizations exhibited significant changes in the auditory cortex. In contrast, IC exhibited a profound and highly stimulus-selective reduction in responsivity following maternal experience (n=243±34 units from N=4-7 expert mice; n=410±44 units from N=6-9 naïve mice). Cohousing significantly reduced the fraction of responsive neurons for five out of six full ultrasonic USVs and the 80 to 40 kHz sweeps (p<0.01 for each). In MGB, for 2 out of 6 multisyllable USVs, proportions of responsive units dropped from ∼30-31% in naive mice to near-zero levels (∼3.3%) in cohoused mice (p*<*0.001 for both; n=30±1 units from N=2-3 expert mice; n=150±12 units from N=6-8 naïve mice). This experience-dependent suppression in IC and MGB was highly specific to multi-syllabic USVs. In contrast, the proportions of neurons responding to isolated syllables were equivalent between naive and alloparental mice (FDR-corrected p>0.05 for all syllables) in all three areas (**Figure 4g**). Together, these results demonstrate that maternal experience drives a targeted, subcortical reduction of population-level responsiveness specifically to full sequences of USV syllables. This selectivity for multi-syllable USVs over single syllables, as well as the supralinear responses we observed in AUD and IC for individual syllables when played in context strongly suggests that these units perform temporal integration over durations of hundreds of milliseconds.

We used principal component analysis (PCA) to examine the population response dynamics to USVs (**Figure 4h-j**). We first pre-processed the data before PCA by pooling all significantly responsive single units across recordings and pseudo-randomly sampling trials of USV playback from all units to create separate high-dimensional state spaces for naive and cohoused mice. trial- averaged responses from pre-stimulus baseline through the duration of each USV onto the first three principal components revealed distinct population trajectories for each USV. Visualizations of the trajectories in dimensions defined by the first three PCs were qualitatively distinct between naïve and expert, with smoother trajectories in the naïve group reflecting more sustained increases in firing rate rather than syllable-locked spikes. Population-level similarity across USV representation, quantified as pairwise trajectory distances for each pair of USVs (**Figure 4j**), was higher in experts compared to naïve. In contrast, trajectory distances did not vary across populations pooled by brain area (**Extended Data Figure 3**). This may reflect a shift from sustained firing rate increases in naïve animals to more temporally sparse population codes in experts, in which overall firing rates are lower but spike timing becomes more aligned with specific USV syllables. As the naïve and alloparent datasets were collected from different mice, the population trajectories lie in distinct state-spaces, and distances between the groups cannot be directly calculated. To quantify temporal differences in trajectories from the two populations independently of cell identities, we calculated Procrustes distance, which identifies rotations of neural manifolds that maximize overlap and thus minimize geometric distance^65^ (**Figure 4k**). To account for inter-animal variability, we assessed statistical significance via a mouse-level permutation test. We found that the population trajectories of naive and cohoused mice were significantly more divergent than expected by chance for the majority of USVs. This indicates that maternal experience drives a fundamental, temporal reorganization of the networks processing infant USVs.

## Maternal experience reduces rate coding of multisyllable USVs in midbrain

Across mice, subpopulations of USV-responsive units exhibited distinct temporal response profiles and selectivity for acoustic stimuli, and the presence of these subpopulations was correlated with alloparenting experience. We examined the relationships between tuning across stimulus features using an unsupervised clustering approach^66^, similar to our approach for comparing USV-responses measured with imaging (**Figure 5**). Peri-stimulus time histograms (PSTHs) were computed for each stimulus and concatenated over time by unit. Concatenated PSTHs were then pooled across mice, omitting non-USV responsive units and recordings in which only a subset of stimuli were played. The pooled population was then projected into a lower dimensional space before clustering (**Figure 5a, Extended Data Fig. 4**). This resulted in five distinct clusters comprised of n=332/422 units (**Figure 5b,c**). All clusters were composed of units across all three areas (**Figure 5a,d**), further supporting a non-hierarchical model of USV representation.

**Figure 5.**
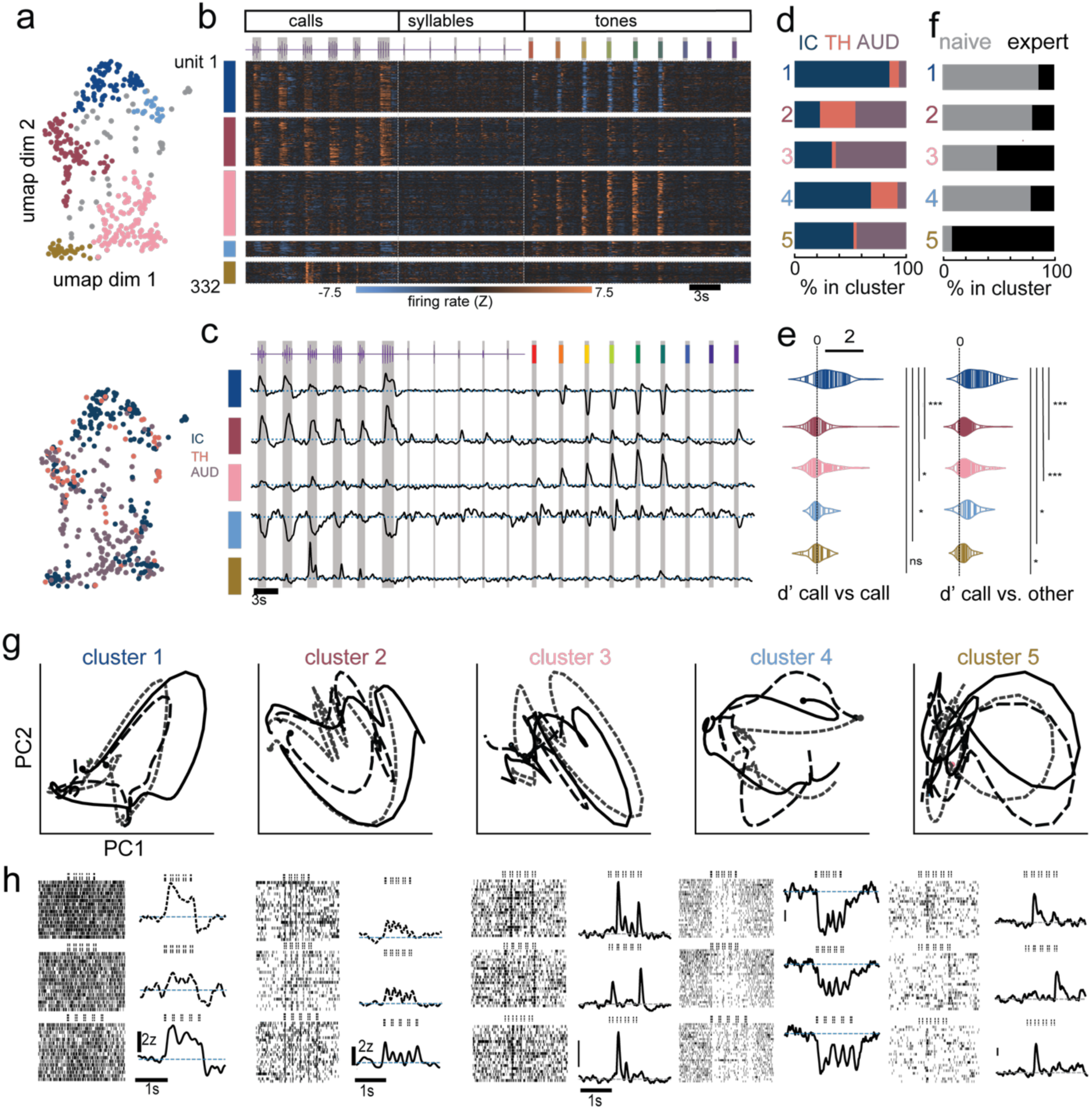
Inter-area functional clusters of neurons respond categorically to pup calls. **a,** Single units in reduced-dimensional space defined by UMAP projection of mean responses of each USV-responsive unit to all acoustic stimuli. Top, overlaid colormap depicting UMAP-based clustering of concatenated PSTHs pooled across recordings. Gray indicates units that were not successfully clustered. Bottom, unit location colormap overlaid on UMAP plot. **b,** Concatenated PSTHs from each clustered single unit sorted by cluster identity. Stimuli were delivered in blocks at time intervals of 3-5s; PSTHs from 1 second prior to onset to 2 seconds post-onset were concatenated by unit for clustering analysis. Heatmap depicts z-scored firing rate relative to 1 second pre-stimulus. **c,** Concatenated PSTHs averaged over single units within clusters. Top, waveforms of concatenated stimuli. Gray vertical bars indicate stimulus timing. **d,** Proportions of units in each cluster from each recorded area. **e,** Left, single unit call vs. call discrimination index (d’) by cluster obtained from Naïve-Bayes classifiers trained on individual units. One-way ANOVA comparing call vs. call dprime across clusters: F=5.8, p=0.0002. Asterisks denote p<0.05, Tukey’s HSD tests for cluster pair comparisons with FDR corrected p-values. Right, d’ for discrimination of calls from all other acoustic stimuli. One-way ANOVA comparing call vs. other d’ across clusters: F=7.4, p=0.00002. Asterisks indicate p<0.05 for individual comparisons. n=332 units. **f,** Percent of units in each cluster recorded in naïve (gray) vs. expert-retrieving (black) mice. P-values obtained by shuffling labels across mice to generate null distributions of proportion permutations. p<0.05 indicates likelihood of true proportions is greater than 95% of all possible permutation likelihoods. **g,** Trial-averaged population trajectories for each cluster’s best 3 USVs projected into the first 2 principal components for visualization for each cluster. Filled circles depict time of trajectory onset. **h,** Single-unit example rasters (left columns) and PSTHs (right columns) for the three best USVs for each cluster in **b**.

Of the three largest clusters, two (clusters 1 and 2, n=93 and n=66 units) were positively modulated by all 6 multi-syllable USVs. Units in cluster 1, of which 85% were in IC, were strongly rate-modulated by USVs, with continuously elevated firing rates from first-syllable onset to last syllable offset (**Figure 5b,c,g,h**). These units were not responsive to individual syllables out of context and were strongly suppressed by pure tone pips in the low-to-middle frequency range (**Figure 5b,c**). To test whether individual acoustic stimuli were categorically encoded in units in each functional cluster, we trained naïve Bayesian decoders on trial-by-trial firing rates in single units, obtaining a discriminability index (d’) for USVs vs other stimuli, as well as USVs from each other (**Figure 5e**). Across clusters, d’ was higher for USV vs. other than for USV vs. USV. Only cluster 1 was strongly discriminative for both measures, with mean d’ >1. Cluster 1 had significantly higher d’ values than all other clusters for USV vs. other, and higher than 3 out of 4 other clusters for USV vs. USV. The strong discriminability of multi-syllable USVs vs other stimuli by cluster 1 is consistent with categorical encoding of pup USVs. Surprisingly, this cluster was strongly biased toward naïve mice that did not retrieve pups (**Figure 5f**), suggesting that encoding of pup USVs in IC is more categorical in naïve virgins than in alloparental mice and could be associated with pup avoidance behavior exhibited by naïve females^10,17^.

In contrast to cluster 1, units in cluster 2 were generally tuned to sound onset, with time-locked responses to individual syllable onsets within calls, syllables played in isolation, and tones (**Figure 5b,c,g,h**). Consistent with non-specific onset responses, mean d’ for both USV vs USV and USV vs other were much lower than in cluster 1. This cluster was also primarily made of up of units from naïve animals. A large cluster (cluster 3, n=111) of primarily auditory cortex units (63% AUD, 4% MGB and 33% IC units) had varied selectivity for USVs and were primarily syllable-selective within USVs (**Figure 5b,c,g,h**), but not significantly modulated by isolated syllables out of context (**Figure 5b,c**). Unlike cluster 1, these units poorly discriminated individual USVs (USV vs USV), but more accurately discriminated USV vs other **(Figure 5e**). Proportions of units in cluster 3 were similar across naïve and alloparent mice **(Figure 5f**), indicating that selectivity for multisyllable USVs is present in all three areas irrespective of parental experience.

## Strengthened IC functional connectivity after maternal experience

Because most clusters contained units from multiple areas, we hypothesized that the distinct tuning properties in each cluster could arise from interarea connections. Therefore we assessed functional connectivity across areas. To identify significant coupling between simultaneously-recorded units independently from shared inputs and stimulus modulation, we modeled spike transmission using fully-coupled population encoding models^67–70^ (**Figure 6, Extended Data Fig. 5**). For each recording, we trained a generalized linear model (GLM) in which the spike train of each unit is predicted by a linear combination of the convolved spike trains of every other simultaneously recorded unit, as well as the timing and identity of each USV (**Figure 6a**). The models assumed that the predicted spike trains followed Poisson statistics and were fit with Group Lasso regularization to impose sparsity and robustly identify significant coupling interactions. After fitting with K-Fold cross validation, we extracted coupling filters describing the probability of spike transmission from one unit to another over a period of 1 second, focusing on auditory cortex and IC due to the abundance of responsive units from those sites. To measure the effect of coupling from each area we fit three separate models with cross validation: (1) a fully coupled model, in which all units were included as predictors in addition to timing of each USV stimulus; (2) an AUD-only model, in which IC units were left out of the feature space, and (3) an IC-only model, in which auditory cortex units were left out of the feature space. For each unit, the contribution of coupling from each area was quantified as the difference in model performance (McFadden’s Pseudo-r^2^) between the fully-coupled model and the left-out model (**Figure 6b-f**).

**Figure 6.**
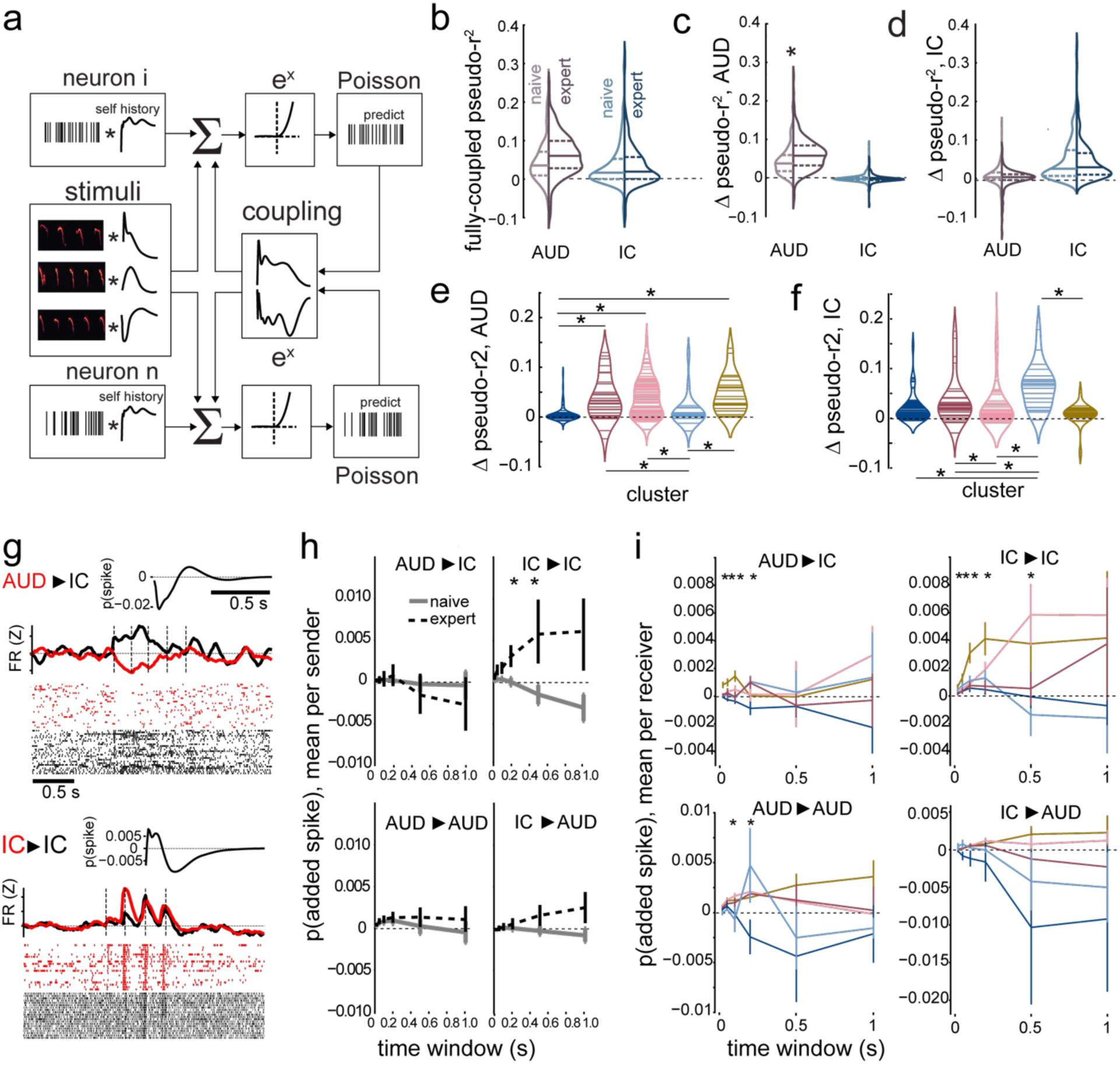
Encoding models of inter-area coupling reveal experience-dependent enhancement of local coupling in IC over timescales relevant for multisyllable integration. **a**, Schematic of fully-coupled encoding models, depicted for two simultaneously recorded example neurons. **b,** Distributions of goodness-of-fit (McFadden’s pseudo-r^2^) for fully-coupled GLMs fit to auditory cortex units (purple) and IC units (blue) during epochs of repeated USV presentation. Darker shades indicate units from expert mice; lighter indicate naïve. **c,** Distributions of per-unit differences in goodness-of-fit between fully coupled model and model in which only auditory cortex units were included as predictors. P = 0.007, Z = 3.12 obtained from GEE test with clustering by mouse comparing naïve vs. expert, AUD units. **d** As in **b**, for difference between full model and IC-only model. **e,** Distributions by cluster of change in model fit (delta McFadden’s pseudo-r^2^) when AUD coupling features are omitted from models. n=249 units, H=57.75, p=9*10^-12^, Kruskal-Wallis Test across clusters. Asterisks indicate p<0.05. **f,** Change in model fit when IC coupling features are omitted. n=259, H=32.68, p=1.4*10^-6^. **g,** Top, paired spike rasters and PSTHs from an auditory cortex unit (red) and a coupled downstream IC unit (black). Top right, spike transmission probability between AUD sender and IC receiver over 1 second, extracted from Poisson Generalized Linear Model (GLM) fit to multi-area single unit spike trains parameterizing pairwise coupling between units in addition to stimulus-driven effects. Bottom, example significantly-coupled IC-IC pair. **h,** Mean spike transmission probability over multiple windows by sender and receiver area. Blue, naïve; pink, cohoused. Asterisks denote p<0.05 for individual time windows, obtained from a Generalized Estimating Equation (GEE) modeling the effect of cohousing clustered by mouse identity and corrected for multiple comparisons using Benjamini-Hochberg False Discovery Rate (FDR) correction. For IC to IC p(spike) within 200 ms: p=0.014, Z=3.45, coef.: 0.007; 500 ms: p=0.019, Z=3.16, coef.: 0.003; n=303 units from N=8 naive mice, n=182 units from N=6 expert mice. **i,** Top left, mean probability of an added spike within 20, 50, 100, 200, 500, and 100 ms of sender unit spike averaged by functional cluster, AUD to IC pairs. (Kruskal-Wallis comparison of net spikes across clusters in 20 ms window: H=25.36, p= 4.3*10^-5^; 50 ms: H=24.35, p=6.8*10^-5^; 100 ms: H=18.04, p=0.001; 200 ms: H=11.21, p=0.024; 500 ms: H=2.08, p=0.72; 1 s: H=2.78, p=0.60; IC units by cluster: cluster 1: n=76, 2: n=13, 3: n=32, 4: n=23, 5: n=10). Asterisks indicate p<0.05, Kruskal-Wallis. Error bars indicate standard error. Top right, IC to IC pairs. (Kruskal-Wallis, IC to IC, 20 ms: H=24.32, p=6.9*10^-5^; 50 ms: H=16.01, p=0.003; 100 ms: H=13.77, p=0.008; 200 ms: H=16.2, p=0.003; 500 ms: H=12.95, p=0.012; 1 s: H=7.47, p=0.11). Lower left, AUD to AUD pairs (Kruskal-Wallis: 100ms: H=10.47, p=0.03; 200 ms: H=10.82, p=0.03; all other windows, p>0.05; AUD units by cluster: 1: n=6, 2: n=18, 3: n=65, 4: n=2, 5: n=14). Lower right, IC to AUD pairs (Kruskal-Wallis: p>0.05, all windows).

Distributions of fully-coupled model performance by unit were equivalent across naïve and experienced mice in both areas (AUD: β= 0.016, p=0.18; IC: β= -0.0003, p=0.91, **Figure 6c**). However, coupling between auditory cortex units was significantly higher in expert mice than in naïve (β=0.018, p=0.002). Auditory cortex coupling was equivalent for IC units across expert and naïve (β= -0.002, p=0.31), and IC coupling was equivalent across expert and naïve in auditory cortex units (β= -0.006, p=0.43) and IC units (β=-0.001, p=0.94, **Figure 6d**). These results suggest that cohousing experience leads to an increase in functional connectivity within auditory cortex during barrages of pup USVs. To determine whether this effect was specific to USVs, we used the same model structures to fit activity during repeated tone presentation (**Extended Data Fig. 5a- c**). Inclusion of cortical coupling had similar effects on model performance in auditory cortex and IC (**Extended Data Fig. 5b)**, as did IC coupling, indicating that the increase in predictive power of intra-cortical coupling was specific to USVs.

Cortical coupling during USV epochs (**Figure 6e,f**) varied substantially across the functional clusters described in **Figure 5**. Strikingly, cortical coupling was lower in cluster 1 than in all clusters other than 4 (cluster 1 vs 2: Z: -4.48, p=0.00002; 1 vs. 3: Z: -6.24, p=4.43*10^-9^; 1 vs. 4: Z: -1.23, p=0.25; 1 vs. 5: Z: -5.56, p=1.37*10^-7^, post-hoc Dunn’s test with FDR correction). Units in cluster 4, which were primarily USV-suppressed units in IC, were also less coupled to auditory cortex that all other clusters except cluster 1. Both clusters 1 and 4 were predominantly present in naïve mice, suggesting that non-cortically coupled units in IC contribute less to USV-modulated ensembles in experts. Conversely, IC to IC coupling was higher in cluster 4 than all other clusters, while cluster 1 was equivalent to all but cluster 4.

We also quantified spike transmission probability between pairs of units over multiple time scales spanning 20-1000 msec to determine the duration over which cohousing altered the strength of functional connectivity between and within AUD and IC (**Figure 6g-i**). For each unit, spike transmission filters were averaged over all downstream units by area to obtain the net impact on each area (**Figure 6h,i; Extended Data Fig. 5h**). Cohousing resulted in a specific pathway-dependent enhancement of spike transmission specifically within the inferior colliculus. This enhancement was timescale-dependent, reaching statistical significance only at windows of 200 ms (β: 0.003, p=0.013) and 500 ms (β: 0.007, p=0.018). These are timescales over which multi- syllabic USVs could be integrated to discriminate temporal structures of calls. In contrast, cortical projections to IC were similar in naïve and maternal animals across all evaluated time windows (all corrected p>0.31). Projections from the IC to auditory cortex were not significantly affected at long timescales (all corrected p>0.09). Surprisingly, given that intracortical coupling was more strongly predictive of cortical activity in experts (**Figure 6c**), spike transmission probability in pairs of cortical units was also unaffected by experience (**Figure 6h, Extended Data Fig. 5e,f**). Our model was not sensitive to dynamics over timescales shorter than 20 ms, which is likely required for inferring monosynaptic spike transmission probabilities within cortex; thus, the increased predictive power of intracortical coupling features in experts may be due to coupling over short timescales.

Although in aggregate, spike transmission from auditory cortex to IC and within cortex were unchanged between naïve and expert, subsets of projections within the naïve and expert groups could be more sensitive to experience. We tested whether spike transmission probability varied across the functional clusters described in **Figure 5**. The probability of added spikes from AUD to IC and IC to IC varied significantly by cluster over most timescales (**Figure 6i**). Within time scales of 20-100 ms, AUD to IC spike transmission was lower in cluster 1 than cluster 5 (20 ms: Z: -4.5, p=4.34*10^-5^; 50 ms: Z: -4.20, p=0.0002; 100 ms: Z: -4.06, p=0.0005; Dunn’s Test with FDR correction) and over 20-50 ms, cluster 1 was lower than cluster 3 (20 ms: Z: -2.42, p=0.038; 50 ms: Z: -2.98, p=0.014). Units in clusters 3 and 5 were primarily present in experts, while cluster 1 was almost entirely units from naïve mice, further supporting a model in which cortico-recipient neurons in IC are more strongly modulated by USVs in experts than in naïve mice. Similarly, AUD to IC spike transmission in cluster 4 was significantly lower than in cluster 5 over 20-100 ms windows (20 ms: Z: -3.05, p=0.011; 50 ms: Z: -2.5, p=0.038; 100 ms: Z: -3.34, p=0.008).

Cluster 5 AUD to IC transmission was also greater than cluster 3 at 20 and 100 ms (20 ms: Z: 2.86, p=0.014; 100 ms: Z: 2.80, p=0.017, and greater than cluster 2 at 100 ms (Z: 2.34, p=0.48). The significant variability in IC to IC spike transmission was primarily driven by lower values in cluster 5 within 20-500 ms. Overall, IC units in cluster 5 were much less coupled to units in both cortex and IC, suggesting that units that are more strongly coupled with both cortical and IC inputs are more critical for USV responses in experts. In contrast to IC, spike transmission probability to auditory cortex units from IC did not vary across clusters (**Figure 6i**, p>0.05 for all time scales, Kruskal-Wallis). Intracortical spike transmission varied significantly by cluster only at 100 and 200 ms windows (**Figure 6**). Post-hoc pairwise comparisons between clusters were only different for pairs including cluster 1, which contained 6 auditory cortex units.

Together, these results support a model in which cohousing experience structurally upregulates correlated spiking dynamics within local IC circuits over long timescales during barrages of USVs, and suppresses activity of IC neurons that are not strongly coupled with cortex or local IC circuits.

## Discussion

These findings indicate a substantial experience-dependent enhancement of temporal integration across multiple syllables of pup distress USVs, coupled with a reduction in the overall excitation elicited in subcortical auditory areas. Because infant cries usually promote avoidance in naïve virgins, the reduction in activity likely represents suppression of circuits that drive avoidance, while the parallel enhancement of temporal integration in subsets of neurons in auditory cortex and IC may reflect a mechanism for increased categorical recognition of distress calls by experienced alloparents.

Many social species use repeated, periodic vocalizations to communicate, and the behavior elicited in conspecifics can rely more on the temporal structure of these calls than on spectral properties of the individual motifs^10,14,15,57^. In mice, USVs elicited by pups, adult males, and adult females can be reliably discriminated based on temporal features despite overlapping frequency ranges^16^, suggesting that the brain could also rely on temporal features to discriminate across categories. The duration of pup distress call inter-syllable intervals (ISIs) is critical for eliciting pup retrieval behavior^10^, and superficial layers of auditory cortex are more sensitive to a broader range of ISIs in maternal females^9,10^. The increased selectivity of CC outputs could arise from spectrotemporal features unique to individual syllables, which are primarily non-harmonic frequency-modulated whistles^71^, or integration over multiple syllables delivered at stereotyped ISIs.

Our multi-site recordings revealed single units in IC, MGB, and cortex each with tightly time- locked spiking elicited by later USV syllables. This indicates that pup call selectivity does not arise simply from hierarchical feature extraction, despite evidence that auditory cortex integrates inputs over longer durations^72^ and is unable to fire synchronously with repeated acoustic stimulus trains at high frequencies, in contrast with MGB and IC^73,74^. The results of our current study also support ISIs as a feature that could be extracted by subcortical neurons in experienced females: in IC, USV-responsive units responded more strongly to later syllables in maternal females. This facilitation indicates integration over syllables, which is necessary for extracting ISI.

In parallel with the increased multi-syllable facilitation in IC in experienced females, we also observed an overall reduction in the proportions of units in IC and MGB that were responsive to multi-syllable USVs. In IC, this reduction was largely due to a decrease in units that exhibited sustained firing rates over the duration of USVs. These units were not selective for later syllables, and were better able to discriminate USVs, as well as USVs vs. other stimuli, than other functionally-clustered units. The presence of these units in naïve animals suggests the existence of a subcortical circuit mediating behavior other than retrieval in response to pup distress calls. Naïve virgins tend to avoid rather than approach pups, and proximity to pups elicits a strong noradrenergic response in locus coeruleus^17,75^ that decreases in magnitude over maternal experience^17^. Additionally, playback of pup USVs alone is aversive to virgins^10^. Thus, sustained- responsive neurons in IC could mediate pup avoidance. Similarly, IC has been found to mediate sound-driven fear and escape behaviors^44^. These units were also less functionally coupled to simultaneously recorded auditory cortex units compared with the other clusters, further supporting the existence of distinct inter-area circuits mediating responses to pup USVs.

Despite the increased magnitudes of delayed and suppressed USV responses we observed in CC neurons, we observed no differences between naïve and experienced animals in overall proportions of neurons that were USV responsive in auditory cortex from our single-unit recordings. This is most likely due to the mixed populations of unlabeled cells we recorded with Neuropixels, in contrast to the projection-specific labeled populations measured with 2-photon calcium imaging and in vivo whole cell/cell-attached recording. Furthermore, tracking labeled CC and CS neurons over days of experience revealed that despite substantial change in individual neuron responses day to day, the overall proportions of labeled neurons that were USV responsive did not change in either projection, consistent with estimates from previous studies^10,54^. Surprisingly, although single-unit responses to multisyllable USVs in auditory cortex were strongly supra-linear compared to isolated single syllables, we did not observe an increase in later-syllable facilitation in experienced animals, as we did in IC. If the experience-dependent increase in delayed USV responses in corticocollicular neurons is due to increased later-syllable facilitation, it is likely that this effect is counterbalanced by a reduction in facilitation in another subset of neurons. Encoding models of single unit spike trains during USV playback revealed that coupling of units within auditory cortex was significantly more predictive in experienced females than naïve animals, and this effect was not observed during blocks of tone stimuli. Thus, it is likely that experience modifies recurrent circuitry within cortex specifically for USV responses.

Cortico-recipient neurons in IC are primarily located in the non-lemniscal shell regions^21,25^, which project densely to lateral/non-lemniscal areas of auditory thalamus including the peri-intralaminar thalamic nucleus (PIL), medial MGB (MGBm), and dorsal MGB (MGBd). These regions integrate multisensory input and have been implicated in a range of naturalistic and social behaviors^76^. A recent study from our lab identified PIL as a key driver of maternal responses to pup USVs in oxytocin neurons of the paraventricular hypothalamic nucleus (PVN), and suppressing activity in PIL impaired pup retrieval^77^. Our finding that corticocollicular projections were necessary for retrieval, as well as the enhanced USV responses we observed in a subset of these neurons suggests that non-lemniscal IC projections to PIL could mediate the transfer of auditory input to this maternal behavior-promoting circuit. In parallel with the reduction in USV responsive units in IC, syllable selectivity via later-syllable facilitation was enhanced in among the smaller population of USV-responsive units in experienced animals. Our findings implicate increased selectivity for temporal features of multisyllabic USVs as a substrate for learning of maternal behavior in a distinct corticofugal circuit, in parallel with suppression of circuits that may drive pup avoidance.

## Acknowledgements

We thank M.A. Kirchgessner, M.A. Long, K.A. Martin, K. Quiñones-Laracuente, A.H. Williams, M. Wollet, and Y. K. Wu for comments and discussions. This work was funded by the BRAIN Initiative (NS107616 to R.C.F., F32 MH123016 and K99DC021581 to A.M.L.), NICHD (HD088411 to R.C.F.), NIDCD (DC12557 to R.C.F.), and NIMH (T32 MH019524 to A.M.L.).

## Methods Animals

All procedures were approved under NYU Grossman School of Medicine Institutional Animal Care and Use Committee (IACUC) protocols, in accordance with National Institutes of Health (NIH) guidelines. Experiments were performed on wild-type virgin female C57BL/6J mice, or in the case of some behavior and chemogenetics assays, RBP4-Cre heterozygous virgin females. For all experiments, mice were between the ages of 5 and 12 weeks old. Virgins were cohoused with wild-type C57BL/6J dams with litters beginning at P0. Timed-pregnant dams were purchased (Taconic, Charles River) and arrived between E10-E17.

## Acoustic stimuli

Acoustic stimuli were delivered via a TDT ultrasonic speaker at a distance of 10 cm from the right ear for all experiments. Stimuli were presented in blocks based on type (e.g., pure tones, full USVs, single USV syllables). Pure tone pips of 500 ms duration ranging from 4-64 kHz at half-octave spacing were calibrated to 70 dB sound pressure level (SPL) and played in pseudo-random order at intervals of 3 or 5 s. Full USV stimuli consisted of 6 multi-syllabic calls recorded from isolated pups aged P0-5. Total durations ranged from 800 ms to 1.2 s, with 4 to 5 individual syllables played per call in the order they were recorded. Calls were chosen to span the typical range of inter-syllable intervals observed in natural pup calls. Sound levels for calls were calibrated to match peak intensity across calls, such that peak intensity was ∼70 dB SPL for each call. Original recordings were edited to remove background noise, and spectrograms were recorded regularly to ensure that no clicks or distortions were played. Individual call syllables from one call were played at 2 second intervals only during Neuropixels recordings.

## 2-photon in vivo calcium imaging

### Cranial window implant and viral transfection surgeries

Survival surgeries were performed using antiseptic surgery technique to place cranial windows over auditory cortex and to inject viral vectors for expression of jGCaMP8f. Briefly, mice were anesthetized with 1.5-2% isoflurane vaporized in oxygen delivered at a rate of 2 liters/minute. Anesthesia was monitored by regularly checking breathing rate and foot-pinch reflex throughout surgery, and mice were kept at an internal temperature of 37° C with a heating pad. Mice were stabilized with a bite bar and nose cone attached to a stereotax. At the onset of surgery buprenorphine (0.1 mg/kg) and dexamethasone (2 mg/kg) were administered intraperitoneally to manage pain and inflammation. Prior to making an incision, the scalp was trimmed of hair and topical lidocaine (2%) was applied. The scalp was cleaned with iodine and 70% ethanol according to proper sterile protocols, and a patch of the scalp was removed to expose the skull. The skull was cleaned of connective tissue and the left temporalis muscle was separated from the skull to expose auditory cortex. Small craniotomies were performed targeting inferior colliculus or posterior striatum using stereotactic coordinates (IC: -5 mm posterior to bregma, 0.8 mm lateral, 0.7 m ventral; PS: -1.7 mm P, 3.4 mm L, 2.5 mm V) and retrograde AAV-Syn-jGCaMP8f-WPRE virus (Addgene) was injected using a Nanoject. Auditory cortex was located using skull and blood vessel landmarks, and a 3 mm area of skull was removed. Dura was left intact, and a circular glass window (FST/Harvard apparatus, thickness 0.1 mm) was applied to the brain surface and fastened to the skull using a combination of cyanoacrylate glue and dental cement. A custom stainless steel headplate was attached to the skull using metabond and all exposed skull was covered. Mice recovered from anesthesia on a heating pad and were individually housed for 24 hours with access to wet food. Additional doses of dexamethasone were administered daily for 2-5 days following surgery to facilitate healing.

### Imaging

Imaging experiments began 3 to 4 weeks after surgery using a 2-photon microscope with resonant galvo scanners and a 20x Nikon objective (Moveable Objective Microscope, Sutter Instruments). Mice were stabilized for imaging by clamping the headplate to a custom holder (Thorlabs parts) and positioning the mouse in a length of acrylic tube. Mice were acclimated to head-fixation daily for several days prior to imaging for increasing durations starting with 5 minutes. GCaMP-expressing neurons were imaged with an excitation wavelength of 900 nm, and epifluorescence was bandpass filtered. 2D fields of CS projecting neurons were imaged at depths of 250-500 µm ventral to the dura, while CC fields ranged from 350-800 µm. Image stacks were collected using ScanImage (Vidrio Technologies) at a rate of 30 frames/second for 512x512 pixel images. Imaging data were collected over multiple days beginning prior to the start of co-housing, with identical stimuli delivered on each imaging day. Auditory stimuli were played with an ultrasonic speaker (TDT) positioned 10 cm from the right ear. In each mouse the same imaging field was imaged each day, using blood vessels and neurons as landmarks to manually match locations, and neuron masks were considered to belong the same neuron on multiple days if the areas across each day were at least 60% overlapping. Data were excluded if changes in window clarity (e.g., from bone re-growth or inflammation under the window) impacted the ability to image previously-imaged neurons. In each mouse, one or two 450 µm^2^ fields were tracked.

### 2-photon imaging analysis pre-processing

Tif stacks were pre-processed using Suite2P. Imaging frames were registered over entire sessions from individual days, then regions of interest (neurons) corresponding to somas were identified semi-automatically using an activity-based algorithm and manual curation. Fluorescence time series were extracted from each neuron over the duration of imaging. Using custom MATLAB scripts, neurons with surrounding neuropil that had much higher fluorescence at baseline were excluded from further analysis. Time series were filtered using a bi- directional median filter with kernel width of 5 frames (each frame ± 2 frames).

### 2-photon imaging analysis

All further analysis was done using custom MATLAB scripts except where noted. dF/F was calculated as (F_t_-F_0_)/F_0_ for each stimulus trial (tone or USV) using a 2 second period prior to stimulus onset as F0. All neurons tracked longitudinally in **Figure 2** were included in analysis. Best early responses to stimuli (**Figs. 1, 1E, 2, 2E**) were defined as follows: for each neuron, responses to each stimulus were averaged over trials, then the mean dF/F was computed over the first 2 seconds following stimulus onset. The “best” USV or tone were defined as the USV and tone evoking the largest magnitude mean dF/F. The same procedure was used to determine the best late response-evoking USV, except that mean dF/F was computed from 2-5 seconds following stimulus onset. For the most suppressed response-evoking stimuli, mean dF/F was computed in the first 3 seconds following stimulus onset, and the greatest magnitude negative dF/F was considered the most suppressed response.

### Longitudinal unit tracking

Neurons were tracked across days by registering Suite2p reference images across pairs of days and computing area overlap of neuron masks using the open source MATLAB program neuronMatchPub (https://github.com/ransona/neuronMatchPub). neurons were considered to be the same neuron if 60% of pixels in the masks two masks were overlapping.

## In vivo whole-cell and juxtacellular recordings

In vivo recordings were performed in N=21 female wild-type mice (3 dams and 18 experienced virgins confirmed to be expert retrievers). One week prior to recording, stainless steel headplates were implanted as described above for cranial window surgeries. A patch of skull covering left auditory cortex was left exposed. For awake recordings, mice were habituated to head fixation daily for three days prior to recording for progressively longer durations, up to one hour the day before recording. On the day of recording, mice were anesthetized with isoflurane (1.5-2% in 2 liter/min oxygen) and a small craniotomy (<2 mm) was performed over auditory cortex. The craniotomy was covered in a layer of sterile saline, then a glass coverslip was temporarily fixed over the craniotomy using kwil-sil. Mice were allowed to recover fully from anesthesia prior to head fixation and recording. For anesthetized recordings, mice were anesthetized prior to craniotomy surgery with an initial dose of ketamine (70 mg/kg) and xylazine (10 mg/kg) delivered via intraperitoneal injection. Additional doses of ketamine were given as needed to maintain anesthesia throughout the craniotomy and recording session (approximately every 30 minutes). Anesthesia level was monitored via tail-pinch reflex and breathing rate measurement.

### Viral injections

For optotagging experiments, 2-3 weeks prior to recording, AAV1-FLEX-ChR2- GFP was injected in auditory cortex as with GCaMP injections) and retro-AAV-Cre was injected in either left posterior striatum or left inferior colliculus using stereotactic coordinates (PS: 3.4 mm ML, -1.7 mm AP relative to bregma and -2.5 mm from brain surface; IC: 0.8 mm Ml, -5.0 mm AP relative to bregma and 0.7 mm from brain surface).

### Whole-cell and cell-attached electrophysiological recordings

Borosilicate glass pipettes (O.D. 1.5 mm, ID 0.86 mm, Sutter Instruments) were pulled into tips of 2.5-5 MΩ resistance. Pipettes were filled with internal solution containing 127 mM K-gluconate, 8 mM KCl, 10 mM phosphocreatine, 10 mM HEPES, 4 mM Mg-ATP, 0.3 mM Na-GTP 527 (osmolality, 285 mOsm; pH 7.2 adjusted with KOH). Tips were mounted on microelectrode holders with AgCl wire. Recordings were obtained using a Multiclamp 700B amplifier and digitized with a 1440B Digidata (Molecular Devices). Data was acquired with Clampex 10.7 (Molecular Devices), low-pass filtered at 1 kHz, high-pass filtered at 100 Hz and digitized at 20 kHz. Pipette tips were lowered 0.5 – 0.8 mm from the pial surface of the brain to target layer 5. Pipette pressure was measured using a manometer (Omega) maintained at 15-20 mbar until the desired depth was reached.

### Optogenetic identification of auditory cortex projection neurons

For identification of light sensitive neurons, whole-cell and cell-attached recordings were obtained using a Fiberoptic Light Stimulating Device with 465 nm blue LED (A-M Systems, Item# 726500) connected to a Fiber Optic Light guide (A-M Systems, Item# 726527). The optic fiber was inserted into the patch pipette via 1.5 mm O.D. Continuous Optopatcher holder (A-M 547 Systems, Item# 663943). Pulses of blue light were delivered through the optic fiber via Digidata 1440A (Molecular Devices) while recording in cell-attached or whole-cell configuration. Light pulses of 20, 50 or 200 ms duration were delivered with increasing intensity from 20 to 100% of full LED power (3 mW at the tip of the fiber). ChR2+ neurons responded to light pulses with an increase firing rate at low jitter <10 ms from light onset, while ChR2- neurons were not modulated by blue light, or if multi-synaptically activated, did not respond with low jitter and high reliability across trials.

### Analysis of in vivo whole-cell and cell-attached recordings

All analysis was performed using custom MATLAB scripts except where noted. Voltage traces for each cell were acquired using Clampex, then imported into MATLAB using the open source program abfload. For spike rate and timing measurements, voltage traces were highpass filtered above 100 Hz and thresholded manually to obtain binary spike trains. Spike time was recorded as the timing of the first threshold crossing. Firing rates were computed using 10 ms-wide bins and averaged over trials of repeated stimuli to obtain PSTHs. Stimulus-evoked firing rates were computed in 2 seconds following stimulus onset and a baseline period averaged over 1.5 seconds was subtracted.

## Behavioral assays with chemogenetics

2-3 weeks following viral injection surgery, virgins were cohoused with a dam and P0-P1 litter of at least four pups in the dam’s home cage. Virgins were co-housed for at least 3 days before retrieval was tested. Prior to test sessions drug administration (CNO or vehicle), virgins were confirmed to be expert retrievers (retrieving pups with at least an 80% success rate). Pup retrieval performance was tested using a trial-based assay as previously described^19,21,22^. Briefly, in each session of retrieval, the test subject was habituated for 45-60 minutes in an arena with corncob bedding, food and hydrogel and 2-3 pups age P0-P3 were placed in a corner of the arena with nesting material from the test subject’s homecage. Habituation and testing took place in a dark room under infrared light and minimal red light to enable the experimenter to perform the assay. During testing, on each trial a single pup was placed in a corner of the cage away from the nest. Trials were initiated only when the test subject was in the nest. Retrieval was scored as successful if the subject brought the pup back to the nest within 2 minutes. If after 2 minutes the pup was not retrieved, the pup was removed from the cage. Video of each session was collected with a USB webcam and ultrasonic audio recorded with an ultrasonic speaker (Avisoft). In a subset of mice, ethograms of behavior were hand-scored using Datavyu and aligned with the timing of USVs, detected using Deepsqueak. For studies of long-term changes with parental experience, one CS- labeled mouse retrieved 100% of pups on day 0 and subsequent days was excluded from analysis.

### Drug administration

Clozapine-2H20 (CNO) was dissolved in sterile saline. 45-50 minutes prior to a retrieval test session, a 2 mg/kg dose of CNO or the equivalent volume of vehicle (saline) was administered via intraperitoneal injection. CNO and vehicle sessions occurred on the same day at least 3 hours apart. The experimenter was blinded to treatment. For several mice, additional sessions were performed the following day with a 1 mg/kg dose of CNO.

## Behavioral assays with optogenetic suppression at presynaptic terminals

For optogenetic manipulations during behavior, pup retrieval assays were performed as described for chemogenetic manipulations. 4-5 weeks prior to behavioral testing, naïve virgins were injected with viral constructs for expression of either eOPN3 (AAV-CaMKIIa-eOPN3-mScarlet-WPRE, addgene) or PdCO (AAV-hSyn-PdCO-eGFP-WPRE, Addgene) in left auditory cortex. During the same surgery, optic fibers were implanted over left IC (400 um diameter, 0.5 NA, trimmed to ∼2 mm length touching the dorsal surface of IC) and left posterior striatum (200 um diameter, 0.39 NA, 5mm length, coordinates as described for viral injections to posterior striatum, 2.5 mm ventral to pial surface). Mice were cohoused with a dam with young pups for at least 3 days prior to testing, and were confirmed to be expert retrievers (retrieving at least 80% of pups when tested with no fiber optic patch cable attached). Mice were habituated to the testing chamber and connection of fiber optic patch cable for at least 1 hour prior to pup retrieval assays. Each mouse was tested on 3 separate blocks: no-light control with optic fiber attached, with light delivered via patch cable to IC, and light delivered via patch cable to posterior striatum. For eOPN3-expressing mice, 0.5s pulses of 525 nm light were delivered every 5s, starting 10 minutes prior to testing and throughout the duration of the assay. For PdCO-expressing mice, 405 nm light was used before and during testing, and following testing, at least 5 repetitions of a 4s pulse of 525 nm light were delivered with an inter-stimulus interval of 10s for inactivation of the construct. Both wavelengths of light were delivered at an intensity of 5 milliwatts. Blocks were spaced in time by at least 30 minutes. Viral expression and optic fiber placement were confirmed via post-hoc histology.

## Head-fixed Neuropixels recordings

Acute Neuropixels silicon probe recordings were performed in 18 wild-type nulliparous female C57BL/6J mice between 6 and 15 weeks old, 7 of which were cohoused with dams and litters (pups <p8) and successfully retrieved pups. Animals were implanted with headplates and habituated as described for in vivo whole-cell recordings. On the day of recording, mice were anesthetized with isoflurane (1-1.5% in 1L/min oxygen) and small craniotomies (<1 mm) were made over auditory cortex and IC on the left hemisphere, localized by stereotactic coordinates. Small durotomies were made to accommodate probe insertion. Craniotomies were covered with duragel (product) followed by kwik-sil (product), and mice were allowed to recover for ∼ 1 hr prior to recording.

Prior to insertion, Neuropixels 1.0 or 2.0 probes were coated with diI or diD (diluted to 1% in etOH) and allowed to dry. Animals were confirmed to be awake and alert, then headfixed for recording. Kwik-sil was removed prior to probe insertion, with duragel left intact. Probe 1 was inserted into Auditory cortex at a slightly negative angle relative to the coronal plane, and orthogonal to the tangential surface of cortex. Probe 2 was inserted into Inferior Colliculus at an angle of 30 degrees relative to the coronal plane and orthogonal to the horizontal and sagittal planes. Both probes were lowered at a speed of ∼3 um/second in the axial dimension using electric micromanipulators (Sutter MP285) until the desired depth was reached (Probe 1: 3.4 mm from the pial surface, Probe 2: 2 mm from the pial surface for 1.0 Neuropixels probe recordings; for 2.0 probes (4 shanks) probes were lowered to 1 mm and 1.5 – 2 mm. Probes were retracted ∼100 um after allowing the brain to settle for 1-2 minutes. Prior to recording, 2% agarose in 0.9M normal saline was warmed to ∼40C and applied to an area covering both craniotomies to improve recording stability and reduce brain pulsation. Recording began after at least 20 minutes or until visible spikes appeared stable across channels.

Acoustic stimuli were delivered through a TDT ultrasonic open-field speaker positioned 10 cm from the right ear. All pure tone stimuli were presented at 70 dB SPL. All USV stimuli, including multisyllable and single syllable USVs, as well as trains of downsweeps were calibrated using Adobe Audition in order to match peak intensity across stimuli (as described previously in Schiavo et al.). Stimuli were played in blocks grouped by type, and were presented in repeating pseudorandom order. Data was acquired continuously during each block.

### Neuropixels data pre-processing and spike sorting

Recording data was acquired using OpenEphys software with a sampling rate of 30 kHz. Raw data stored in the OpenEphys binary format was pre-processed using the SpikeInterface python library as follows: first, AP band streams (for Neuropixels 1.0 probes) or full spectrum data (2.0 probes) were concatenated sequentially across recording blocks into continuous streams for each channel. The raw traces were then high-pass filtered with a minimum frequency cutoff of 400 Hz to eliminate low-frequency fluctuations and local field potentials. Noisy or inactive channels were automatically detected and removed from the recording matrix, and the remaining channels were phase-shifted to account for the temporal delays associated with the sequential sampling of the Neuropixels probe channels. A common median reference (CMR) was calculated globally across all valid channels and subtracted from each channel to minimize correlated electrical noise across the probe. We used the DREDGE algorithm within SpikeInterface to correct for probe drift and longitudinal tissue movement over the duration of the recording. Motion was estimated using a spatial sliding window approach, with both the window step size and window scale set to 100 µm. Recording blocks were excluded from further analysis if substantial motion was detected. Following pre-processing and motion correction, the corrected AP traces were routed to Kilosort4 for automated spike sorting. Because probe drift and motion correction were already compensated for during the SpikeInterface pre- processing stage, the internal drift correction built into Kilosort4 was disabled. Clusters found by Kilosort4 were then manually curated in Phy2 and clusters confirmed to be single units were labeled and included in further analysis.

Following spike sorting, the sorting outputs were loaded into SpikeInterface to perform comprehensive quality control and feature extraction. For each unit, dense waveforms were extracted over a window from 1.3 ms before to 2.6 ms after the spike peak, from which unit templates (average, median, and standard deviation) were computed. Quality metrics and features were calculated for each unit, and units were subsequently curated, and only those classified as ’good’ were retained for downstream analysis. The concatenated, continuous spike times for all units across both probes for each recording were stored, along with metadata for each unit, in objects constructed with the pynapple (Python Neural Analysis Package) python package. All further analysis was performed on these reduced data objects.

### Post-hoc histological localization of single units

To anatomically localize recorded neurons, spike sorting outputs were integrated with stereotactic coordinate frameworks using histological reconstructions. Following recordings, brains were sectioned and imaged to visualize the fluorescent DiI/DiD tracks left by the Neuropixels probes.

To determine the precise anatomical trajectory of each probe, these histological slice images were processed using the MATLAB-based SHARP-Track library. Using SHARP-Track, the histological slices were downsampled and non-linearly morphed to register them to the Allen Mouse Brain Common Coordinate Framework (CCFv3) coronal atlas. The fluorescent probe tracks were manually marked on the registered slices, allowing for the reconstruction of the 3D probe trajectory and the generation of a reference table mapping physical distance along the probe to specific Allen CCF brain areas and stereotactic coordinates. The anatomical location of each recorded unit was subsequently determined by mapping the relative depth of the unit on the Neuropixels probe (extracted from the Kilosort sorting output) to the SHARP-Track generated structural annotations using custom Python scripts. This process yielded an estimated 3D stereotactic coordinate (Anterior-Posterior, Medial-Lateral, Dorsal-Ventral relative to Bregma) and a brain area label for each isolated unit, allowing downstream analyses to explicitly control for anatomical location.

## Neuropixels data analysis

### significance testing for stimulus responsiveness

To robustly detect statistically significant stimulus-evoked neural responses for each acoustic stimulus, the Zenith of Event-based Time- locked Anomalies (ZETA) test was applied to the un-binned spike trains^78^. The ZETA test evaluated whether the temporal distribution of spikes within a target window (-1000 ms to +2000 ms relative to stimulus onset) significantly deviated from a stationary baseline rate. For units demonstrating a significant response (p < 0.05), the direction of the response (increase or decrease) was determined by comparing the mean firing rates calculated across matched pre-stimulus and post-stimulus evaluation windows.

### Proportions responsive

To determine if cohousing alters the broad probability that a neuron will respond to auditory stimuli, we quantified the fraction of significantly responsive units across experimental groups and brain areas, and modeled the effect using hierarchical statistics. For each recorded neuron, responsiveness to a given stimulus (e.g., specific calls, syllables, downsweeps, and pure tones) was determined using the previously calculated ZETA-test scores. To compare the fraction of responsive units between naive and expert retrieving mice, we modeled the binary responsiveness variable using Generalized Estimating Equations (GEE) implemented via the statsmodels library in Python. Because the dependent variable was binary (responsive vs. non- responsive), a Binomial family link function was used. Crucially, to avoid the statistical inflation associated with pseudo-replication (treating hundreds of neurons from the same animal as independent samples), the GEE model was structured to cluster observations by biological replicate, defining mouse as the random grouping variable. The main effect of the binary predictor variable (cohoused) was extracted from the model, yielding a raw p-value representing the probability that the observed difference in responsiveness arose by chance.

Because responsiveness was tested independently across a large array of distinct stimuli, a multiple comparisons correction was necessary. Within each brain area independently (e.g., AUD, IC), the raw GEE *p*-values across all evaluated stimuli were strictly corrected using the Benjamini- Hochberg False Discovery Rate (FDR) procedure. A corrected FDR p-value <0.05 was considered statistically significant.

### Syllable adaptation index

To determine whether evoked firing rate during multi-syllable USVs exhibited adaptation or facilitation for later syllables compared to the first in a sequence, we first computed baseline mean-subtracted firing rate from the start of each syllable to the beginning of the following syllable for each unit (or up to 200 ms following onset for the last syllable). For each USV-responsive unit, we identified the multisyllable USV evoking the greatest change in firing rate, and within that USV, identified the “best syllable” as the syllable occurring most closely prior to the PSTH peak. We then calculated adaptation index for each unit as the raw best syllable firing rate – firing rate to the first syllable, divided by the sum of the two syllables’ firing rates.

### Population PCA, Procrustes disparity

Neural population dynamics were analyzed by aggregating single-unit responses across multiple mice. To ensure robust representations, analysis was restricted to multi-syllable USV responsive units. We excluded recordings in which fewer than 6 multi-syllable USVs were presented. Because recordings yielded varying numbers of trials per stimulus across different mice, a permutation-based subsampling approach was implemented. For each stimulus condition, the minimum number of available trials across all individual mouse sessions was determined. Trials from each mouse were then randomly subsampled without replacement to match this minimum count, guaranteeing that all mice contributed equally to the pooled pseudo-population tensors, thereby preventing mice with higher trial yields from biasing the population trajectory.

To visualize and quantify population-level neural dynamics, Principal Component Analysis (PCA) was applied to the firing rate data. For each unit, trial-averaged peristimulus time histograms (PSTHs) were computed for each stimulus. These mean PSTHs were baseline-subtracted and Z-scored independently for each unit to normalize baseline variations and equalize the contribution of high- and low-firing neurons. Standard PCA was then fitted to the standardized, trial-averaged population responses to define an orthogonal basis space capturing the directions of maximal variance across time and stimuli. To evaluate trial-to-trial variability, individual single- trial responses were subsequently standard-normalized and projected through the pre-fitted PCA axes, producing single-trial trajectories within the shared population subspace.

To quantify experience-dependent divergence in neural representations, the geometric similarity of population trajectories between naive and expert retriever groups was evaluated using Procrustes analysis. Because naive and cohoused populations consisted of different distinct neurons, they occupy different neural state spaces. To compare them, independent PCAs were first run on the naive and cohoused populations separately. The cumulative explained variance was evaluated for each group to determine the number of principal components required to capture at least 80% of the response variance. The maximum of these two values was selected as a standardized dimensionality to ensure both groups were evaluated in a sufficiently descriptive, dimensionality-matched subspace. The mean trajectory for each stimulus was projected into this matched *N*-dimensional space. Procrustes alignment was then performed to optimally rotate, scale, and translate the naive trajectory to match the cohoused trajectory. The residual error from this alignment (Procrustes disparity) provided a quantitative metric of the fundamental geometric divergence between the two population representations.

To assess the statistical significance of the observed Procrustes disparities, a hierarchical permutation testing framework was utilized. Importantly, to avoid false positives arising from pseudo-replication, permutations were performed at the mouse level rather than at the level of individual units or trials. For every possible permutation of the labels (271 permutations total), the expert/naive status of the mice was shuffled while preserving the true experimental ratio of naive to cohoused animals. For each permutation, the exact same analytical pipeline was executed from scratch: pseudo-naive and pseudo-cohoused populations were assembled, independent PCAs were fitted, data was projected into the dimensionality-matched subspace, and the Procrustes disparity was recalculated. This generated a robust null distribution of geometric disparities expected by chance given the inter-mouse variability. The final statistical significance for each stimulus was calculated as the proportion of the null distribution permutations that yielded a disparity greater than or equal to the true experimentally observed disparity.

### UMAP/hDBSCAN clustering

To characterize the temporal dynamics of neural responses across different auditory stimuli, trial-averaged peristimulus time histograms (PSTHs) were estimated for each unit. Analysis was restricted to all 6 multi-syllable USV exemplars, 5 individual syllables from one of the exemplars, and pure tones (4 kHz to 64 kHz at quarter-octave spacing). We excluded recordings in which any of these stimuli were not presented from this analysis. For each stimulus condition, a continuous PSTH was calculated over a standardized window spanning from 1 second before to 2 seconds after the stimulus onset, using a bin width of 40 ms. These stimulus-specific PSTHs were then concatenated chronologically along the time axis to create a single, continuous, high-dimensional response vector (a "concatenated PSTH") representing each unit’s comprehensive response profile across the entire stimulus set. Only USV-responsive units localized to auditory cortex (AUD), inferior colliculus (IC) or medial geniculate body (MGB) were included in clustering analysis. Units were considered USV responsive if they had a statistically significant change in firing rate following at least one exemplar USV (zeta statistic >2 / p<0.05). In addition, recordings were excluded if any recording blocks with the relevant stimuli were excluded due to recording drift.

To facilitate the identification of distinct functional response phenotypes, the concatenated PSTHs were first temporally smoothed using a 1D Gaussian filter (sigma = 1 bin). The smoothed response vectors were then Z-scored independently for each unit across the concatenated time series to emphasize differences in temporal response shape rather than absolute firing rate magnitudes. Because the concatenated PSTHs represent a high-dimensional feature space, Uniform Manifold Approximation and Projection (UMAP) was employed for dimensionality reduction. Finally, unsupervised clustering was performed directly within the lower-dimensional UMAP embedding space. Hierarchical Density-Based Spatial Clustering of Applications with Noise (HDBSCAN) was utilized to identify discrete functional clusters. The parameters for UMAP (number of nearest neighbors) and HDBSCAN (minimum number of samples, minimum cluster size) were selected following a parameter sweep evaluating cluster stability and noise fraction, and were ultimately selected to minimize the number of unclustered units, as well as to minimize the number of resulting clusters to the smallest number that adequately partitioned obvious variation across groups of units without over-splitting. The selected parameters were number of nearest neighbors = 12, minimum number of samples = 5, minimum cluster size = 20.

### Single unit decoders

To decode stimulus identity from single-unit activity, single-trial spike trains were first converted into continuous firing rate estimates. For each unit and stimulus presentation, spike times within a -1 to +2 second window relative to stimulus onset were extracted. To capture smooth temporal dynamics on a single-trial basis, Kernel Density Estimation (KDE) was applied to the spike times using a Gaussian kernel with a bandwidth of 20 ms. This yielded a continuous trial-by-time feature matrix for each unit. Units with insufficient spiking activity (fewer than 50 spikes) or zero-variance temporal features were excluded from the decoding analysis. To prevent overfitting and reduce the high dimensionality of the temporal response profiles, Principal Component Analysis (PCA) was employed^79^. For each unit, the trial-by-time density matrix was standard-normalized (Z-scored across trials for each time bin). PCA was then applied to extract the most prominent temporal features, reducing the data to a maximum of 5 principal components using a randomized SVD solver. For each unit, a Gaussian Naive Bayes classifier was implemented in scikit-learn to predict the stimulus identity from the PCA-reduced single-trial responses. Each classifier was trained and evaluated using Stratified K-Fold cross-validation (up to 5 folds, dynamically reduced for stimuli with fewer trials) to ensure robust, out-of-sample performance estimates.

Classifier performance was quantified by computing a symmetric discriminability index (d’) for every possible pair of stimuli. A cross-validated confusion matrix was generated by aggregating predictions across all test folds. For any given pair of stimuli, the hit and false alarm rate were extracted from the confusion matrix. To prevent infinite values, probabilities were clipped with a small epsilon (10^-6). The pairwise d’ was then calculated as the difference between the inverse cumulative normal distributions of the hit and false alarm rates (Z(Hit Rate) - Z(False Alarm Rate)). This calculation was performed symmetrically for both directions and averaged, providing a robust metric of how discriminable the neural representations of any two stimuli were for a given unit.

### Generalized Linear Models (GLM) for functional connectivity inference

To model neural firing rates and infer functional connectivity between AUD and IC while accounting for stimulus-driven activity, a Poisson Generalized Linear Model (GLM) was implemented independently for each recording using the nemos (JAX-based Neural Modeling and Simulation) python library. Spike trains for each unit were counted in 20 ms bins. The GLM was constructed using two primary classes of predictors, each convolved with a set of temporal basis functions to capture extended temporal dynamics without overfitting. To model stimulus-driven responses, stimulus event times were convolved with a raised cosine log-stretched basis. The basis set consisted of 6 basis functions spanning a 1.5-second temporal window following stimulus onset. To model functional connectivity, the binned spike counts of all other simultaneously recorded units within AUD and IC were used as predictors. These coupling histories were modeled using an extended raised cosine log basis consisting of 20 basis functions spanning a 1-second temporal window. Because the inclusion of all simultaneously recorded neurons as predictors results in a highly parameterized model, Group Lasso regularization was employed. Group Lasso encourages sparsity at the group level; in this case, driving all temporal basis weights connecting a specific pre-synaptic neuron to a post-synaptic neuron to exactly zero if the connection does not meaningfully improve model fit. The optimal regularization strength (λ) was selected empirically. A parameter sweep was performed across a logarithmically spaced range of λ values. For each lambda, a 5-fold chronological cross-validation was conducted. The models were standardized prior to fitting, and the optimal lambda was chosen as the value that maximized the median out- of-sample McFadden’s pseudo-r^2^ across the cross-validation folds.

Three distinct GLM architectures were fitted for comparison: a "Fully Coupled" model incorporating stimulus history, AUD coupling, and IC coupling; an "AUD Only" model that excluded IC units as predictors, and an "IC Only" model that excluded AUD units as predictors.

Following the full-dataset fit using the optimized *_λ_*, the learned basis coefficients were transformed back into the time domain to yield continuous temporal filters. For inter-neuronal coupling filters, specific physiological metrics were extracted, including peak/trough amplitudes, peak/trough latencies, and the area under the curve (AUC).

Following GLM fitting, the model outputs (feature importance, coupling filters, and cross-validated predictive performance for each model architecture) were aggregated across mice and stimulus sets (2 blocks of 3 calls and one block of tones) for quantitative evaluation. To evaluate the overall efficacy of the functional coupling models, the out-of-sample McFadden’s pseudo-r^2^ was extracted for the Full model (stimulus + AUD + IC), the AUD-only coupling model, and the IC-only coupling model. The unique predictive contribution of cross-area versus local coupling was isolated by computing the difference in r^2^ between nested models. For example, the specific contribution of AUD network activity to an IC neuron’s firing rate was defined as the difference in pseudo-r^2^ between the Full model and the IC-only model.

### GLM Multi-Window Connectivity and Transmission Probability Methods

To dissect the temporal dynamics of inter-areal and local functional connectivity, post-hoc analyses were performed on the time-domain coupling filters extracted from the Generalized Linear Models (GLMs). The raw GLM coupling filters represent the instantaneous modulation of a receiver neuron’s firing rate by a sender neuron’s spike. To quantify the cumulative impact of these connections, the Area Under the Curve (AUC) of the coupling filters was computed using trapezoidal integration. To evaluate how functional connectivity evolves over different integration timescales, the filters were sliced and integrated across multiple progressively expanding temporal windows: 20 ms, 50 ms, 100 ms, 200 ms, 500 ms, and 1 second. To translate the AUC metric into a physiologically interpretable value, the AUC for each pair was multiplied by the baseline firing rate of the receiver neuron. This yielded a transmission probability representing the absolute number of extra (or suppressed) spikes fired by the receiver neuron per individual spike from the sender neuron within that specific integration window.

Because GLM coupling filters can take positive or negative values, the transmission probabilities were explicitly parsed to evaluate pathway-specific Excitatory/Inhibitory (E/I) balance. Putative excitatory connections were defined as those generating a net positive transmission probability (transmission probability > 0). Putative inhibitory connections were defined as those generating a net negative transmission probability (transition probability < 0). The total excitatory drive, total suppressing drive, net spikes, and ratios were independently aggregated across the AUD◊AUD, AUD◊IC, IC◊AUD, and IC◊IC pathways to assess how experience reshaped specific directional motifs over varying temporal windows.

To quantify density of functional networks, a binary ‘significant connection’ threshold was established. A sender neuron was considered to significantly drive a receiver if the absolute transmission probability met a minimum threshold of 1%. The fraction of significant senders out of all possible senders within a specific brain area was calculated to define the connection probability for each receiver neuron. To rigorously compare continuous connection strengths and binomial connection probabilities between naive and cohoused mice, Generalized Estimating Equations (GEEs) were utilized via the statsmodels python library: continuous connection strengths (transmission probability) were modeled using a Gaussian GEE, and connection probabilities (fractions of significant senders) were modeled using a Binomial GEE, weighted by the total number of recorded sender neurons. Both GEE architectures defined mouse identity as the grouping variable. This correctly accounted for within-subject correlation and prevented the statistical inflation (pseudo-replication) that occurs when treating hundreds of neuronal pairs from the same mouse as independent biological samples. Finally, to control for the large number of comparisons across multiple temporal windows and pathways, all raw p-values were strictly corrected using the Benjamini-Hochberg False Discovery Rate (FDR) procedure (α=0.05).

---

**Supplementary Figure 1.**
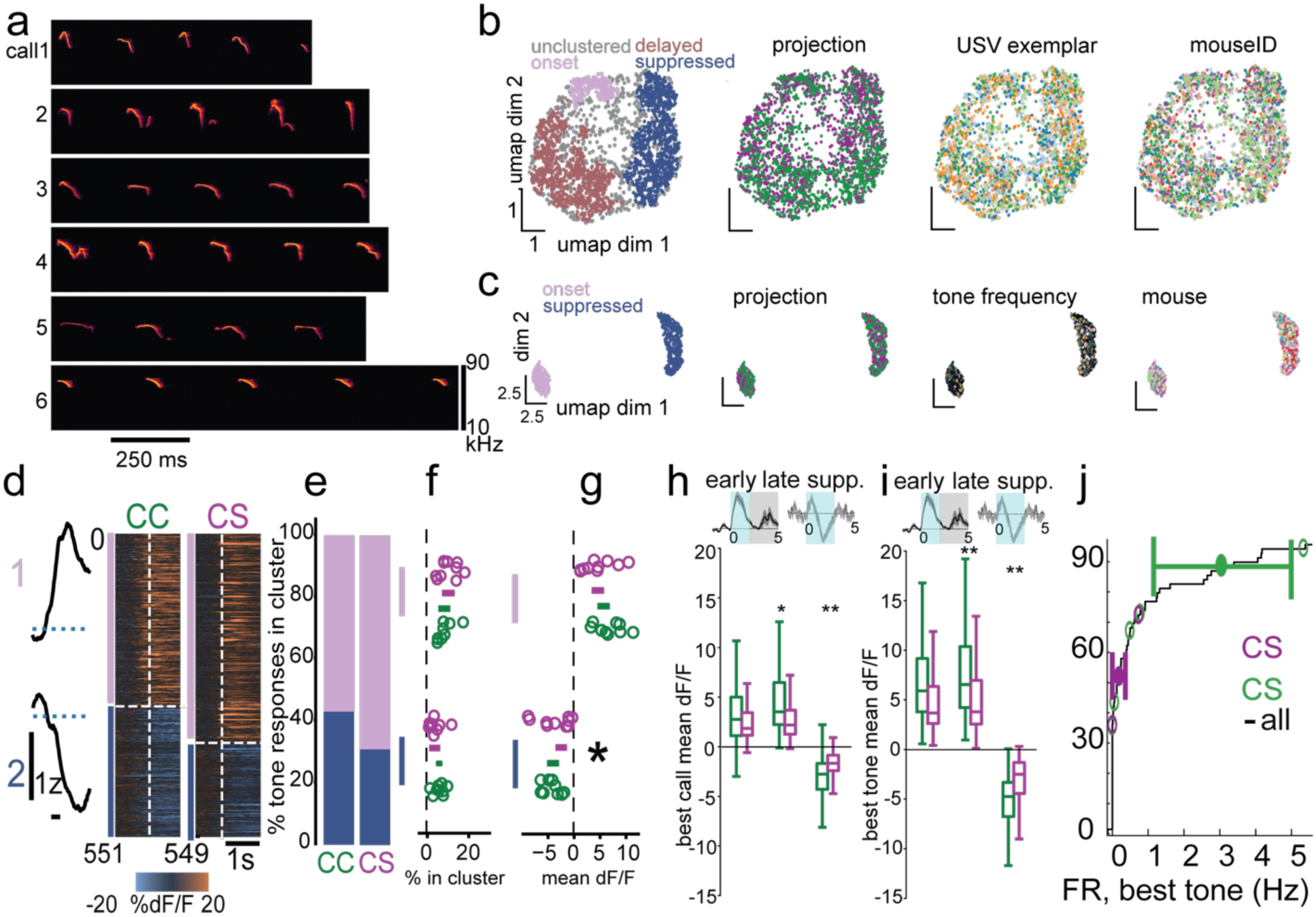
Analysis of neural and behavioral responses to pup calls. **a**, Spectrograms of multisyllable USVs used as stimuli for in vivo imaging and electrophysiology. **b,** USV responses pooled across recordings plotted in reduced-dimension UMAP space based on trial-averaged dF/F during USV presentation. Each point denotes one response from one neuron. From left, units are color-coded by HDBSCAN cluster: projection, USV exemplar, and mouse. **c,** UMAP/HDBSCAN computed on significant tone responses. **d-g,** Clustered trial-averaged significant tone responses. **d**, Left, means of all significant evoked responses in each cluster; right, trial-averaged evoked responses sorted by cluster; **e**, significant evoked responses in each cluster from CC and CS neurons pooled across mice. **f**, percentages of responses in each cluster for each individual imaging field. Circles indicate fields; horizontal bars indicate mean +/- SE pooled by projection. (Mixed effects linear model with SE clustered by imaging field; β: 0.025, p=0.16, CC vs CS proportions in excited cluster; β: 0.022, p=0.21, CC vs CS proportions in suppressed cluster; β: 0.04, p=0.09, projection x cluster). **g,** Mean dF/F of tone responses in each cluster, averaged within imaging field. **h**, Mean dF/F evoked by each neuron’s best early-facilitated (left), late- facilitated (middle), and most suppressed (right) USVs, pooled across mice by projection. **, p<0.01; only least squares linear models with cluster-robust standard errors grouped by mouse. **i**, Mean dF/F for best tone responses. **j**, Firing rates to best tone measured from in vivo cell-attached and current clamp recordings.

**Supplementary Figure 2.**
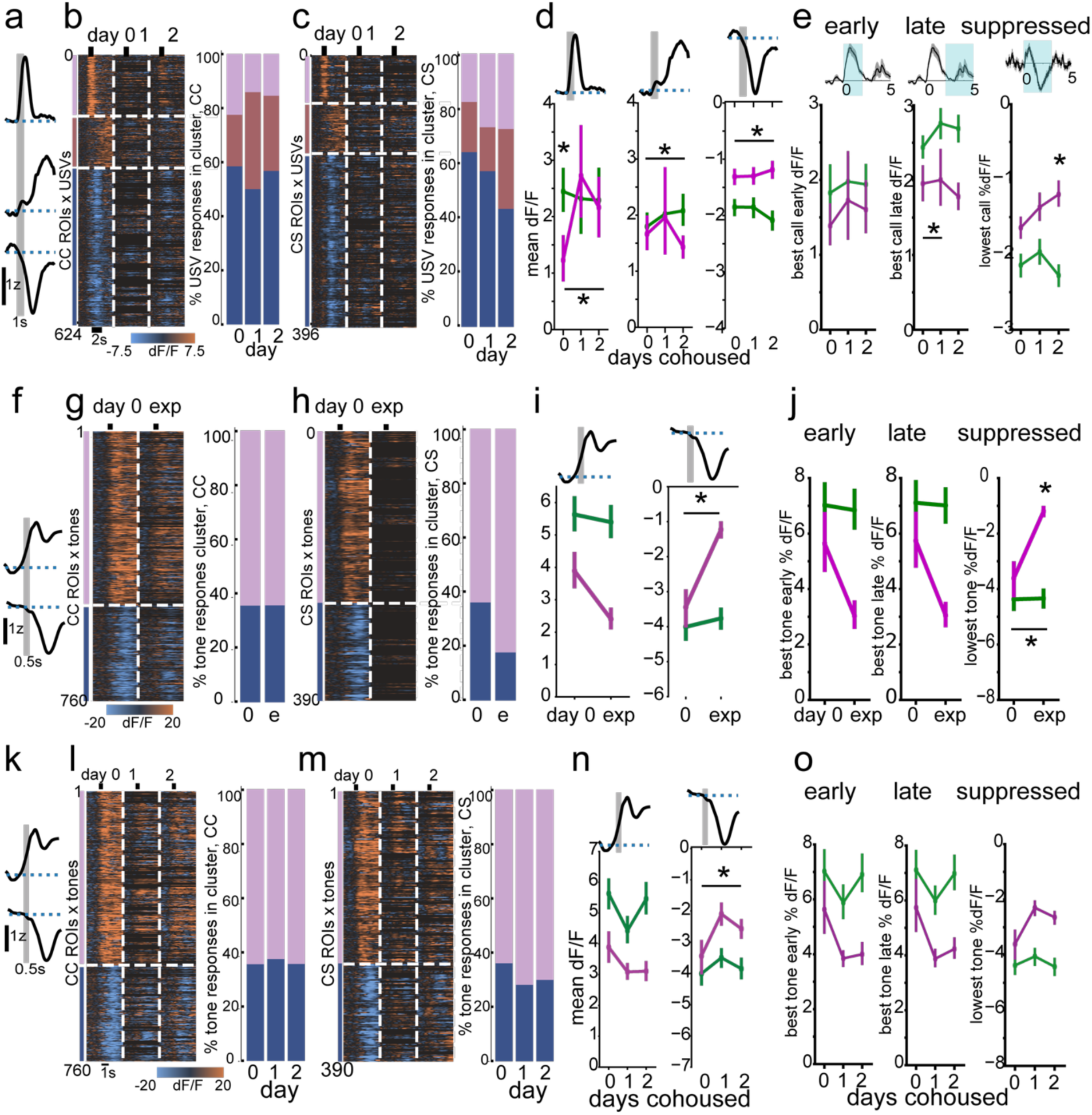
**Daily USV and tone responses in CC and CS neurons. a-d**, Clustered trial-averaged responses to USVs over consecutive days of cohousing. **e**, Best early, late, and suppressed responses to USVs. **f-i,** Clustered trial-averaged responses to tones on day 0 of cohousing (day 0) and first day of expert behavior (exp, e). **j,** Best early, late, and suppressed tone- evoked mean dF/F for each USV-responsive neuron’s best tone, pooled across mice within CC and CS groups on day 0 and first day of expert behavior. **k-o**, Tone responses over days of cohousing. *, p<0.05. P-values obtained by fitting only-least-squares linear regression models SE clustered by imaging field to account for within-field correlation across individual neurons.

**Supplementary Figure 3.**
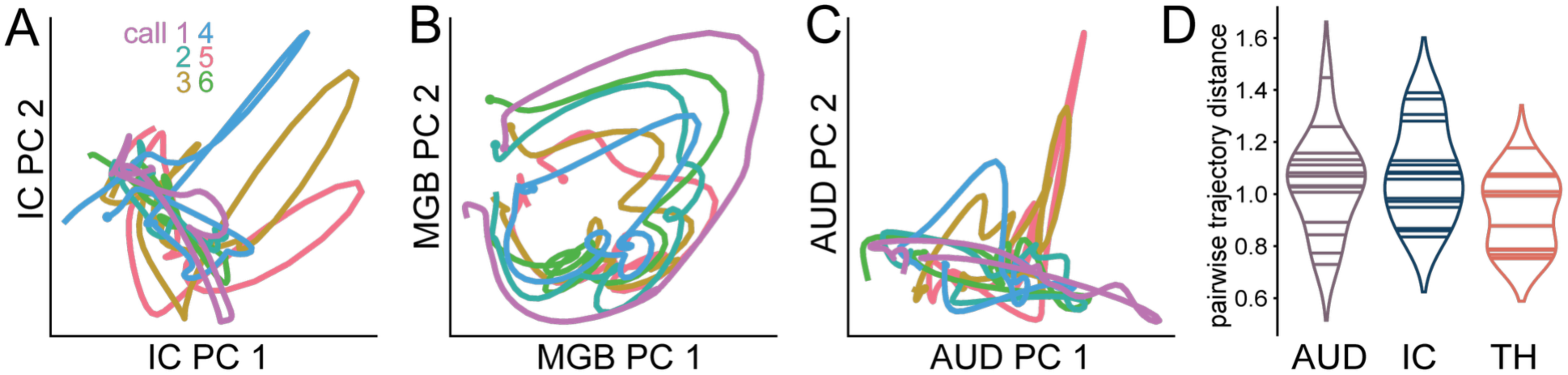
Neural population responses to USVs by brain area. a, Trial- averaged population trajectories for each USV projected into the first 2 principal components for visualization for all USV-responsive neurons in IC, pooled across mice. Filled circles depict time of trajectory onset. **b,c,** As described in **a**, for MGB and auditory cortex. **d,** Distributions of pairwise USV trajectory distances by area.

**Supplementary Figure 4.**
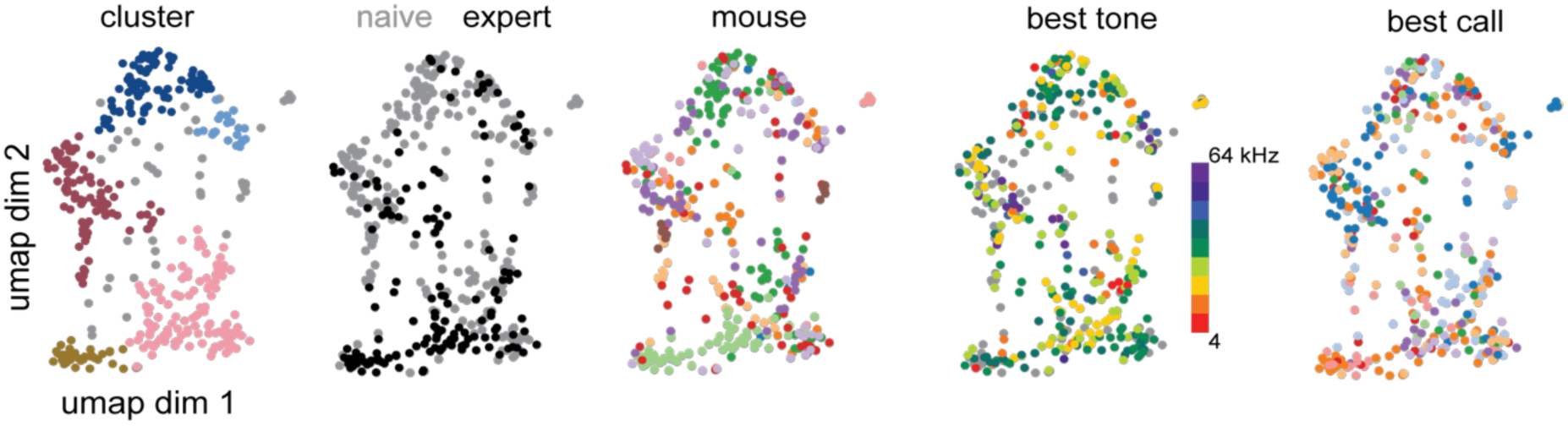
Single unit metadata in UMAP space. Single units in reduced- dimensional space defined by UMAP projection of mean responses of each USV-responsive unit to all acoustic stimuli, color-coded from left to right by cluster, experience, mouse identity, best tone, and best call.

**Supplementary Figure 5.**
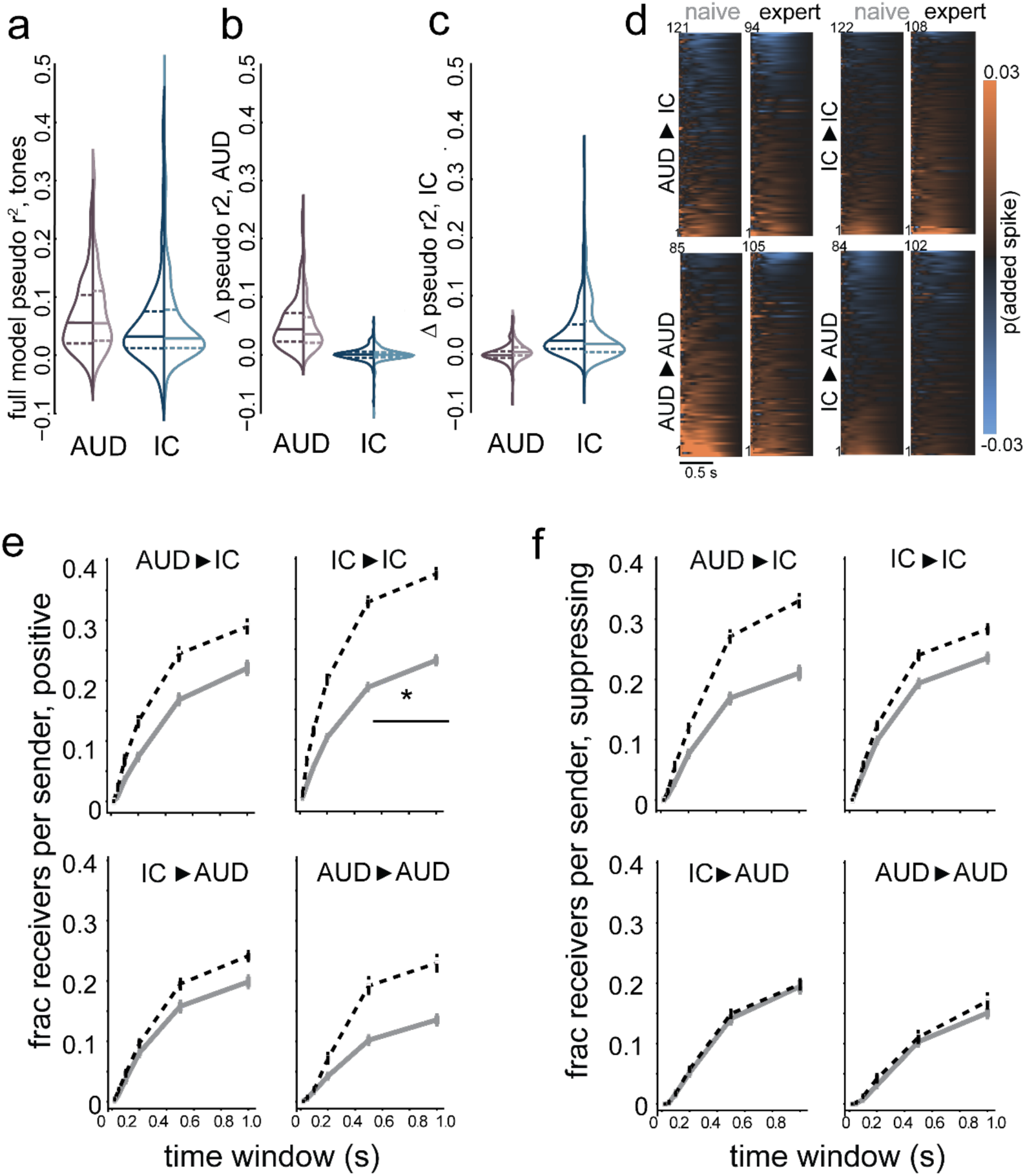
a-c. As described for Figure 5 b-d, for models fit to spike trains during delivery of pure tone stimuli. **d,** Mean spike-transmission probability between each significantly- driving sender unit and all receiver units, averaged by receiver area, for naïve and expert. Only sender-receiver pairs with at least a 1% spike transmission probability over 500 ms were included in averages. Top left, AUD senders to IC receivers; bottom left, AUD to AUD; top right, IC to IC; bottom right, IC to AUD. Each row depicts mean by area for one sender unit. **e,** Mean fraction of significantly excited receivers per sender, by sender and receiver areas, over multiple time windows. IC to IC, 500 ms: p = 0.046, Z = 2.87 obtained from binomial GEE with clustering by mouse comparing naïve and cohoused, FDR corrected. 1 s: p = 0.046, Z = 2.83. **f,** As in E, for significantly inhibited receivers.

